# The fungal peptide toxin candidalysin induces distinct membrane repair mechanisms compared to bacterial pore-forming toxins

**DOI:** 10.1101/2025.05.09.653080

**Authors:** Roshan Thapa, Victor Kayejo, Claire M. Lyon, Bernhard Hube, Julian R. Naglik, Peter A. Keyel

## Abstract

The common fungal pathogen, *Candida albicans*, relies on the pore-forming toxin candidalysin to damage host cells. Cells counteract pore-forming toxins by Ca^2+^-dependent mechanisms, such as microvesicle shedding and annexin recruitment to resist cholesterol-dependent cytolysins like streptolysin O (SLO), or annexin involvement and patch repair in the case of aerolysin. However, the specific Ca^2+^-dependent repair pathways engaged in response to candidalysin remain poorly understood. Here, we determined the involvement of different Ca^2+^-dependent repair mechanisms to candidalysin and compared responses to SLO and aerolysin using flow cytometry and high-resolution microscopy. We report that candidalysin triggered Ca^2+^-dependent repair, but patch repair and ceramide failed to provide significant protection. MEK-dependent repair and annexins A1, A2 and A6 contributed partially to repairing damage caused by candidalysin. However, annexin translocation after candidalysin challenge was delayed compared to SLO or aerolysin challenge. Surprisingly, extracellular Cl^−^ improved cell survival after candidalysin challenge, but not after challenge with SLO or aerolysin. Finally, we found that candidalysin is removed via extracellular vesicle shedding. These findings reveal that Ca^2+^-dependent microvesicle shedding protects cells from candidalysin and can be engaged by multiple molecular mechanisms during membrane repair.

**Graphical Abstract.**
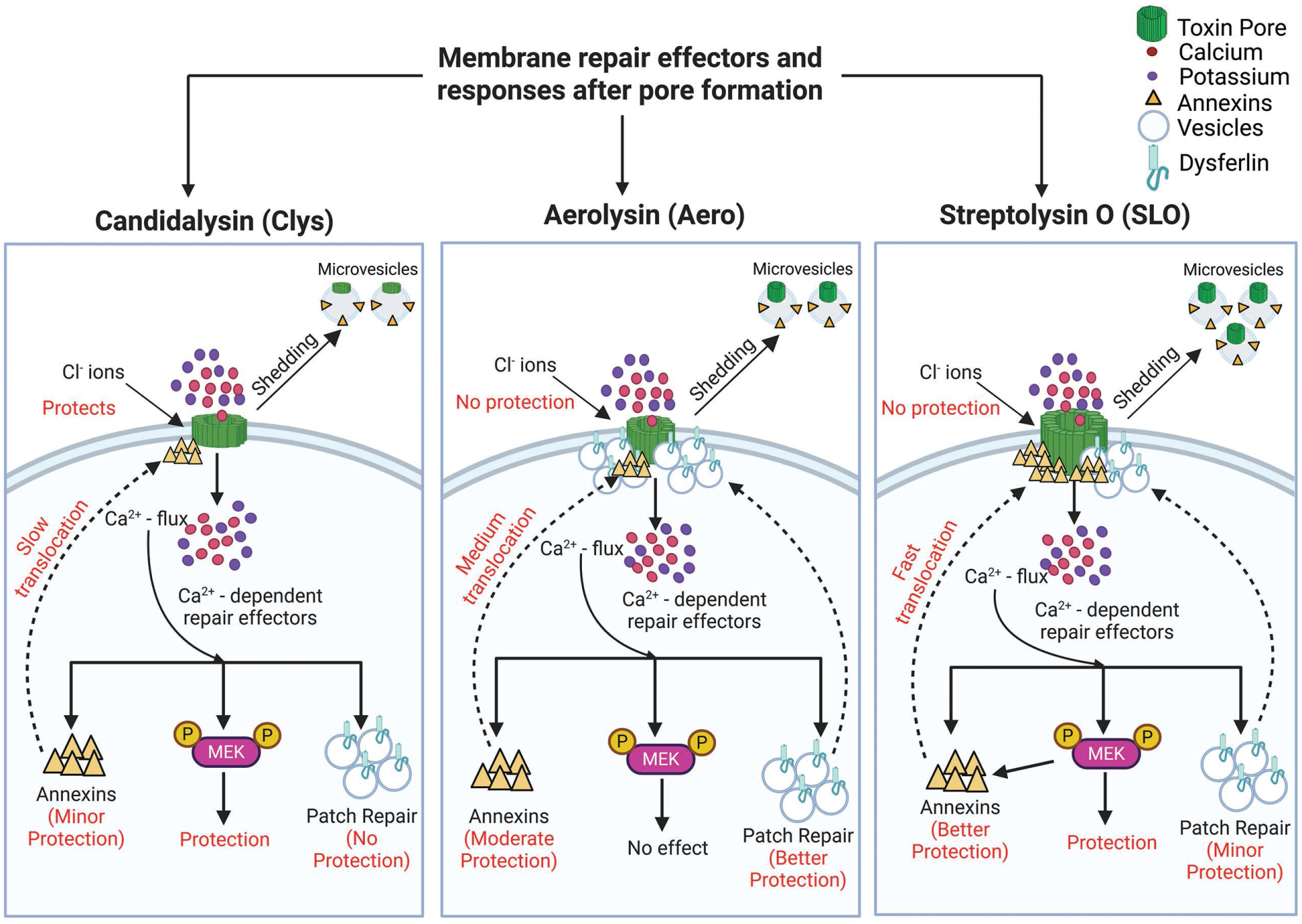
Candidalysin is resisted by distinct repair mechanisms compared to bacterial PFTs. After pore formation and membrane damage by each toxin, multiple repair pathways are triggered downstream of Ca²⁺ flux. Candidalysin induces a protective Cl^−^ influx and activates MEK-dependent repair, which contributes to cell protection. Annexin translocation occurs slowly and provides minor protection, while patch repair is ineffective. In contrast, aerolysin does not benefit from Cl^−^ influx or MEK protection. Aerolysin triggers moderate annexin translocation and relies primarily on patch repair as the main protective mechanism. Streptolysin O elicits rapid annexin translocation and activates MEK signaling, both of which contribute to robust protection. Patch repair plays only a minor protective role against SLO. The figure was created using BioRender.

## Introduction

One leading cause of mucosal and life-threatening systemic fungal infections is *Candida albicans*^1^. The morphological transition of *C. albicans* from a yeast to a filamentous hyphal form is crucial for host cell invasion^2^. Host cell damage after hyphal invasion is promoted by secretion of a key virulence factor—the peptide toxin candidalysin^3^. The precursor peptide of candidalysin is embedded in the polyprotein Extent of cell elongation 1 protein (Ece1p) ^3^. Candidalysin is released from Ece1 by Kex1p and Kex2p mediated proteolytic processing to produce Ece1-III_62–93_ ^3–5^. Candidalysin directly interacts with host plasma membranes to form lytic pores that cause epithelial damage and cell death^3, 4, 6^. Candidalysin may act by binding to several surface components, including glucosaminoglycans, GP1b-α and CD11b, that may promote lytic activity indirectly or activate cell signaling responses^7–10^. While host damage by bacterial toxins has been extensively studied, the mechanisms by which cells survive candidalysin attack are less explored.

Several Ca^2+^-dependent membrane repair mechanisms protect cells from damage. Pathways that resist pore-forming toxins (PFTs) include the physical blockade of the pore (clogging), autophagy, the hetero/homotypic fusion of internal vesicles (patch repair), and removal of the toxin on microvesicles (microvesicle shedding) ^11, 12^. Mechanistically, a wide variety of proteins and lipids enable these pathways to reseal damage. Clogging of pores occurs when annexins form crystalline arrays on the membrane^13, 14^. Autophagy can reseal damage by recruitment of microtubule-associated protein 1A/1B-light chain 3 (LC-3) and formation of the autophagosomal membrane^15^. Patch repair represents the fusion of endosomes and lysosomes with the plasma membrane, which then reseals pores via vertex fusion^12^. A wide range of proteins mediate patch repair, including the muscle protein dysferlin^12^. Toxin pores can be removed via sequestration and shedding on microvesicles^16–19^. Shedding can be energy and protein-independent, but lipid-dependent^16, 19^, or activated by proteins. Proteins that drive shedding during repair include the endosomal sorting complex required for transport (ESCRT) ^18^ and mitogen activated protein kinase kinase (MEK) signaling, partially via annexin A2^17^. Other annexins may also be shed on microvesicles^17, 20^. However, different pore-forming toxins trigger distinct subsets of these repair pathways^21^. Understanding which mechanisms are triggered by candidalysin will help identify key pathways that promote repair.

Prior work with candidalysin investigating repair responses identified three repair pathways. Repair against candidalysin is Ca^2+^-dependent, and can involve ESCRT-III mediated blebbing, lysosomal exocytosis and autophagy^3, 15, 22, 23^. Candidalysin also triggers MEK signaling that promotes epithelial cell survival at late time points^24^. However, the role of other Ca^2+^-dependent repair mechanisms in resisting candidalysin damage, and the relative importance of each pathway, remains unknown.

A comparative analysis of PFTs is expected to reveal which pathways are best targeted to improve cellular resistance to specific toxins. Our prior work compared two other PFTs: streptolysin O (SLO) from *Streptococcus pyogenes* with aerolysin from *Aeromonas hydrophila*^21^. SLO and aerolysin belong to two distinct, well-studied PFT subfamilies, the cholesterol dependent cytolysins (CDCs) and the aerolysin family, respectively^25^. These toxins are repaired via distinct mechanisms^21^. About 70% of the Ca^2+^-dependent repair for SLO uses MEK-dependent microvesicle shedding, while annexins and ceramide elevation also promote survival^17, 21^. In contrast, Ca^2+^ overload kills aerolysin-challenged cells, while patch repair promotes resistance to aerolysin^21^. Therefore, we selected SLO and aerolysin to compare the relative importance of repair mechanisms to resist candidalysin.

Here, we compared the mechanisms of repair between candidalysin and two bacterial pore-forming toxins that engage distinct repair mechanisms. We found that candidalysin was not resisted via patch repair or ceramide elevation. We observed partial protection from MEK-dependent repair and annexins A1, A2 and A6, though annexin translocation after candidalysin challenge was delayed. Surprisingly, we found that elevating extracellular Cl^−^ improved cell survival against candidalysin, which was not observed for SLO or aerolysin. Finally, microvesicle shedding removed candidalysin from cells. Overall, these data suggest candidalysin triggers a distinct subset of membrane repair mechanisms.

## Results

### Candidalysin cytotoxicity against nucleated cells is intermediate between SLO and aerolysin

To compare repair mechanisms between candidalysin, SLO and aerolysin, we first normalized toxin activity using hemolytic activity in human erythrocytes (Supplementary Fig S1). We used hemolytic activity instead of toxin mass because it normalizes toxin activity for better comparison, provides a measure of cellular resistance mechanisms, is consistent with our prior work, and controls for variation in the specific activity between toxin preparations^17, 21^. We challenged HeLa cells with a range of toxin doses, calculated the specific lysis due to toxin, fit the dose-response curves to a logistic model, and calculated the toxin dose needed to kill 50% of the cells (LC_50_) for each experiment^26^ (Fig 1). We first determined the minimum time range to best measure cell damage and repair for each toxin to minimize non-specific lysis. We challenged HeLa cells with candidalysin, aerolysin and SLO for 30 min, 1 h and 2 h and measured the cell viability by MTT and LDH assay (Supplementary Figs S2, S3). Similar to aerolysin, cytolytic activity by candidalysin plateaued at 1 h, whereas SLO plateaued at 30 min (Supplementary Fig S2). Therefore, we used 1 h for candidalysin and aerolysin and 30 min for SLO to compare repair mechanisms.

**Figure 1.**
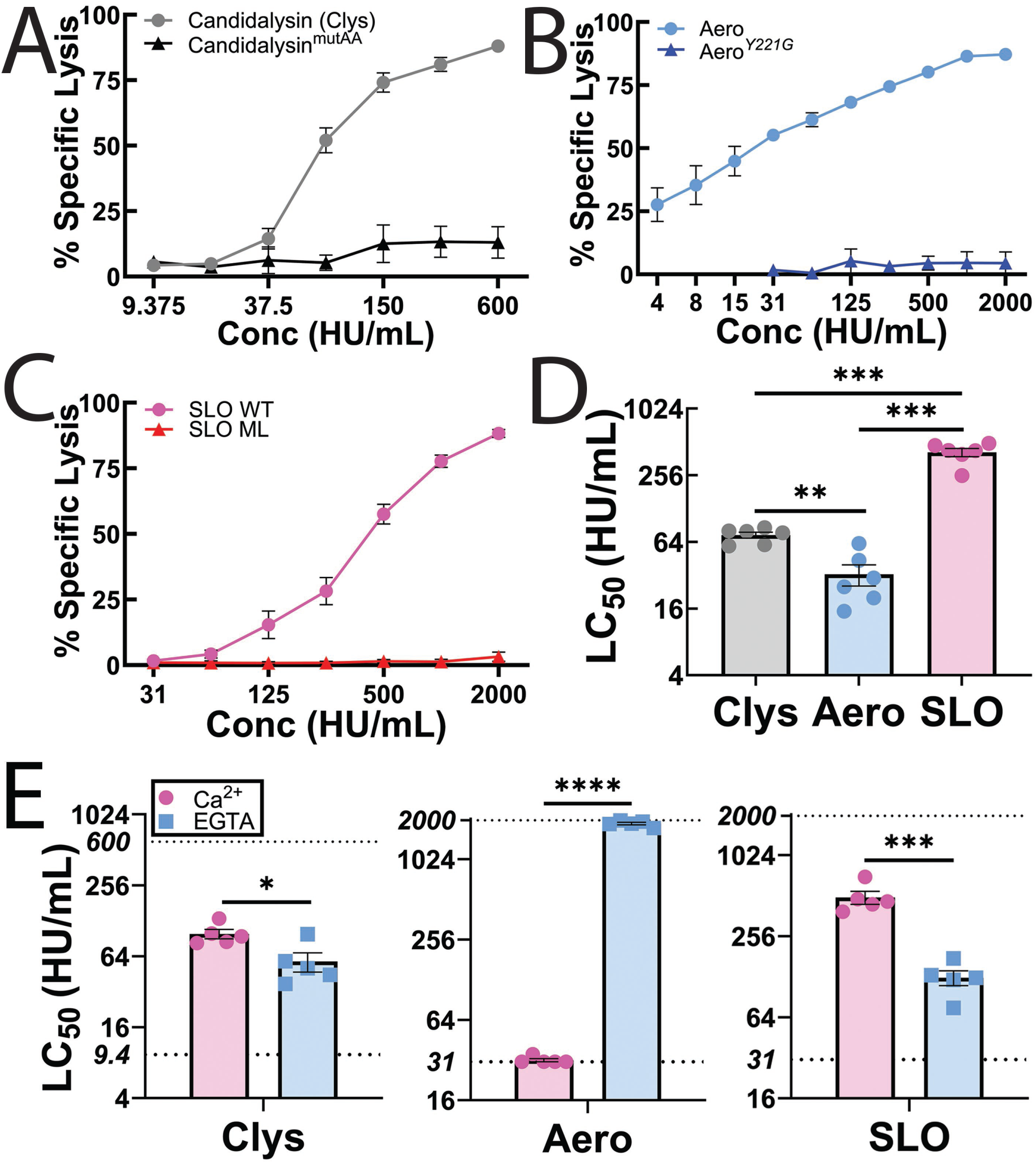
Candidalysin killing is intermediate between aerolysin and CDCs. HeLa cells were challenged with (A) 9.375-600 HU/mL candidalysin (Clys), (B) 4-2000 HU/mL aerolysin (Aero), (C) 31-2000 HU/mL SLO, or (A-C) a mass equivalent to wild-type toxin of candidalysin^mutAA^, aerolysin^Y221G^ , or monomer-locked SLO (SLO ML) in RPMI supplemented with 2 mM CaCl_2_ and 20 μg/ml propidium iodide (PI) at 37°C. PI uptake was analyzed by flow cytometry and % specific lysis was determined. (D) The amount of toxin needed to kill 50% of the cells (LC_50_) was calculated by logistic modeling. (E) HeLa cells were challenged with 9.375-600 HU/mL Clys, 31-2000 HU/mL Aero, or 31-2000 HU/mL SLO at 37°C in RPMI supplemented with either 2 mM CaCl_2_, or 2 mM EGTA. PI uptake was analyzed by flow cytometry. The LC_50_ was calculated as described above. Dotted lines indicate the lower and upper limits of detection. Graphs show the mean ± S.E.M. of (A-D) six or (E) five independent experiments, with (D, E) independent experiments plotted as data points. * p <0.05, ** p<0.01, *** p<0.001, **** p<0.0001 by repeated-measures one-way ANOVA with Tukey’s multiple comparison test.

We next controlled for cytotoxicity due to any impurities in toxin purification by comparing wild-type toxins with an equivalent mass of toxins mutated to cripple cytolytic activity using flow cytometry. Mutation of two key lysines to alanines in candidalysin (candidalysin^AA^) reduced its hemolytic activity by 90% (Supplementary Table S1). Despite residual hemolytic activity, none of the defective toxins were cytotoxic to HeLa cells (Fig 1A-C). In contrast, wild-type candidalysin was less toxic than aerolysin, but more toxic than SLO (Fig 1). The LC_50_ for candidalysin (73.7 ±4.65 HU/mL) was between aerolysin (32.6 ±7.08 HU/mL) and SLO (426.1 ± 38.8 HU/mL) (Fig 1D). These data suggest that candidalysin triggers less repair than SLO, but more than aerolysin.

### Candidalysin triggers Ca^2+^-dependent repair

Most membrane repair mechanisms described to date require Ca^2+^ influx. One key difference between aerolysin and SLO is that Ca^2+^ influx is toxic following aerolysin challenge, but protective against SLO^21^. Previous results showed that Ca^2+^ influx is protective during candidalysin challenge^3, 22, 23^. We confirmed the impact of Ca^2+^ influx on cell survival following candidalysin challenge by chelating extracellular Ca^2+^ with EGTA. Chelating Ca^2+^ in the extracellular medium sensitized HeLa cells to candidalysin, similar to what we observed for SLO (Fig 1E, Supplementary Fig S4A). In contrast, removal of extracellular Ca^2+^ protected HeLa cells from aerolysin (Fig 1E, Supplementary Fig S4A), consistent with previous results^21^. These data suggest that candidalysin engages Ca^2+^-dependent repair mechanisms, similar to those triggered by SLO.

### Patch repair and ceramide fail to protect cells from candidalysin

Our prior work^21^ suggested that patch repair is a backup repair mechanism for SLO, while the primary mechanism for aerolysin. Since candidalysin and SLO both trigger protective Ca^2+^ influx, we hypothesized that patch repair would be a minor contributor to repair. To test this hypothesis, we measured the ability of the well-characterized muscle patch repair protein dysferlin^12, 21^ to promote survival against candidalysin. Dysferlin-deficient C2C12 myoblasts were more sensitive than control C2C12 myoblasts to aerolysin and SLO (Fig. 2A, Supplementary Fig S4B), consistent with prior results^21^. However, loss of dysferlin did not increase cellular sensitivity to candidalysin (Fig. 2A, Supplementary Fig S4B). These data suggest that the patch repair protein dysferlin does not contribute to repair triggered by candidalysin.

**Figure 2.**
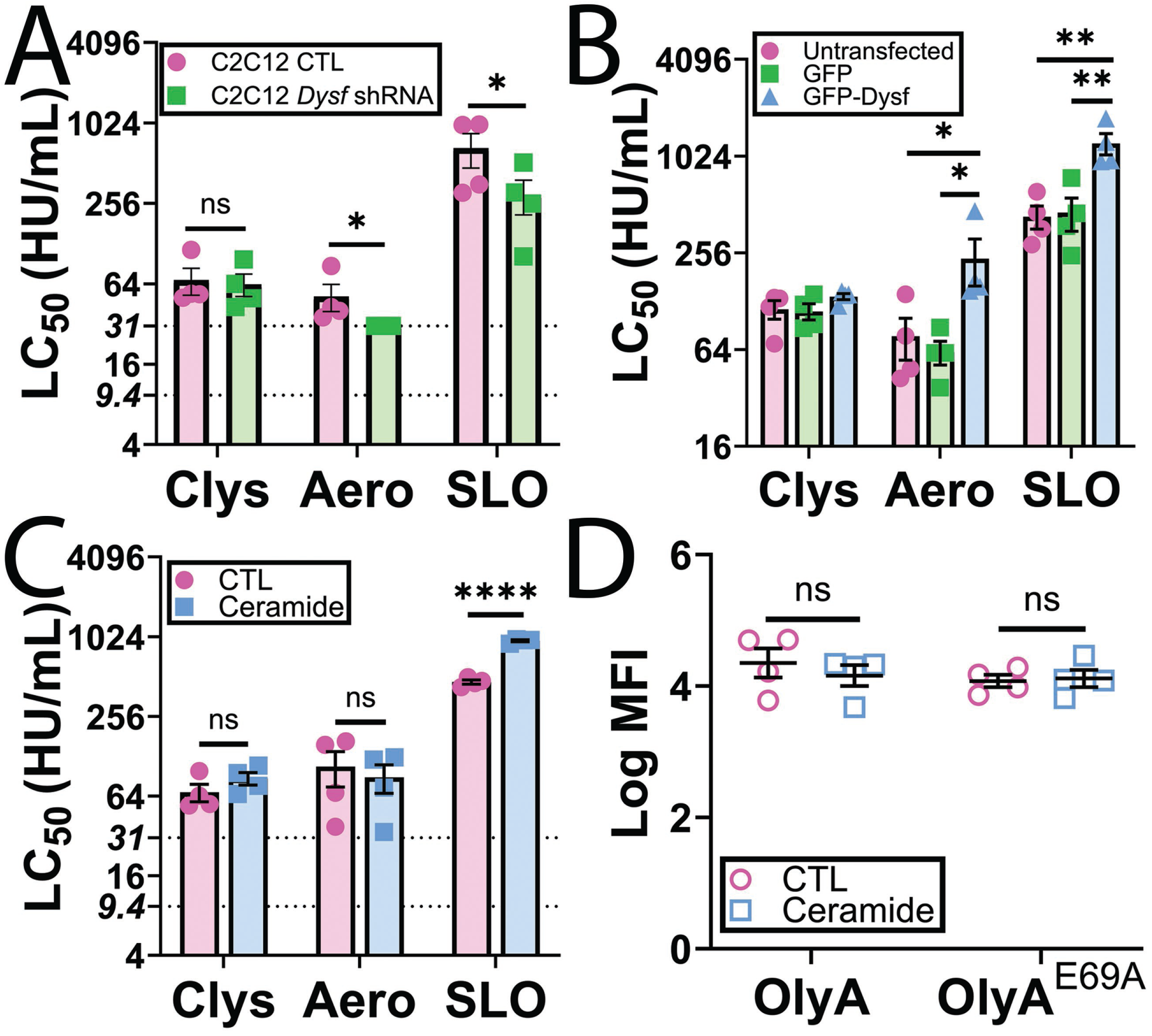
Patch repair and ceramide fail to protect cells against candidalysin. (A) C2C12 cells stably expressing control (CTL) or *Dysf* shRNA were challenged with 9.375-600 HU/mL candidalysin (Clys), 31-2000 HU/mL aerolysin (Aero), or 31-2000 HU/mL SLO. (B) HeLa cells were untransfected or transfected with GFP or GFP-Dysferlin (GFP Dysf) for 48 h and then challenged with 9.375-600 HU/mL Clys, 31-2000 HU/mL Aero, or 31-2000 HU/mL SLO. (C, D) HeLa cells were pretreated with either vehicle DMSO (CTL) or 40 μM PPMP for 72 h. Cells were then (C) challenged with 9.375-600 HU/mL Clys, 31-2000 HU/mL Aero, or 31-2000 HU/mL SLO or (D) labeled with 20 μg/mL OlyA-mCherry WT or OlyA ^E69A^-mCherry for 30 min. PI uptake or OlyA-mCherry binding was analyzed by flow cytometry. The LC_50_ was calculated as described in the methods. The dotted lines indicate the lower limits of detection (9.4 HU/mL for Clys, 31.25 HU/mL for Aero and SLO). Graphs show the mean ± S.E.M. of four independent experiments. * p <0.05, ** p <0.01, **** p<0.0001, ns not significant by repeated-measures one-way ANOVA with Tukey’s multiple comparison test.

To confirm that patch repair is not significant for protection against candidalysin, we measured the ability of dysferlin overexpression to protect non-muscle cells from cytotoxicity. Our prior results showed that over-expression of dysferlin protects HeLa cells from both aerolysin and SLO^21^. Consistent with prior results^21^, we found that GFP-dysferlin transfection enhanced cellular resistance against aerolysin and SLO compared to GFP-transfected cells(Fig 2B, Supplementary Fig S4C). In contrast, GFP-dysferlin transfection failed to enhance cellular resistance to candidalysin (Fig 2B, Supplementary Fig S4C). Based on these data, we conclude that dysferlin-mediated patch repair fails to protect cells from candidalysin.

Since patch repair failed to protect cells against candidalysin, we next tested other mechanisms of repair. Sphingolipids and ceramide contribute to protection against CDCs in multiple systems^17, 27–29^. We previously showed that ceramide accumulation by inhibiting UDP-glucose ceramide glucosyltransferase protects cells from CDCs without perturbing sterol or sphingomyelin levels^17^. We tested the hypothesis that ceramide accumulation also promotes cellular resistance to candidalysin. While ceramide accumulation protected cells from SLO as previously described^17^, it did not alter cell sensitivity to either aerolysin or candidalysin (Fig 2C, Supplementary Fig S4D). To confirm that we did not alter cellular levels of either sphingomyelin-cholesterol complexes, nor total sphingomyelin, we measured them using wild-type ostreolysin A (OlyA)–mCherry^30^, or OlyA^E69A^-mCherry^31^, respectively. Ceramide accumulation did not alter these lipids in the plasma membrane (Fig 2D). Based on these findings, we conclude ceramide does not provide significant cellular protection against candidalysin.

### Chloride ions protect cells from candidalysin

We next tested the contribution of K^+^ efflux to candidalysin resistance, because prior results^32^ suggested it contributes to resisting pore-forming toxins. We blocked K^+^ efflux by adding 150 mM KCl to the culture medium^33–35^. While blocking K^+^ efflux did not impact cell survival following SLO or aerolysin challenge, blocking K^+^ efflux protected HeLa cells from candidalysin (Fig 3A, Supplementary Fig S5A). We next determined the impact of extracellular KCl in a second cell type, primary bone marrow-derived macrophages (BMDM). We used caspase1/11^−/-^ BMDMs to rule out toxin-induced pyroptosis^36^. KCl protected BMDM from both candidalysin and aerolysin, but sensitized cells to SLO (Fig 3B, Supplementary Fig S5B). To determine whether Ca^2+^ influx or K^+^ efflux were more important, we blocked both, and measured cell viability after toxin challenge. The presence of KCl overcame any EGTA-induced sensitivity to candidalysin in BMDM (Fig 3B, Supplementary Fig S5B). Based on these results, extracellular KCl protects cells against candidalysin.

**Figure 3.**
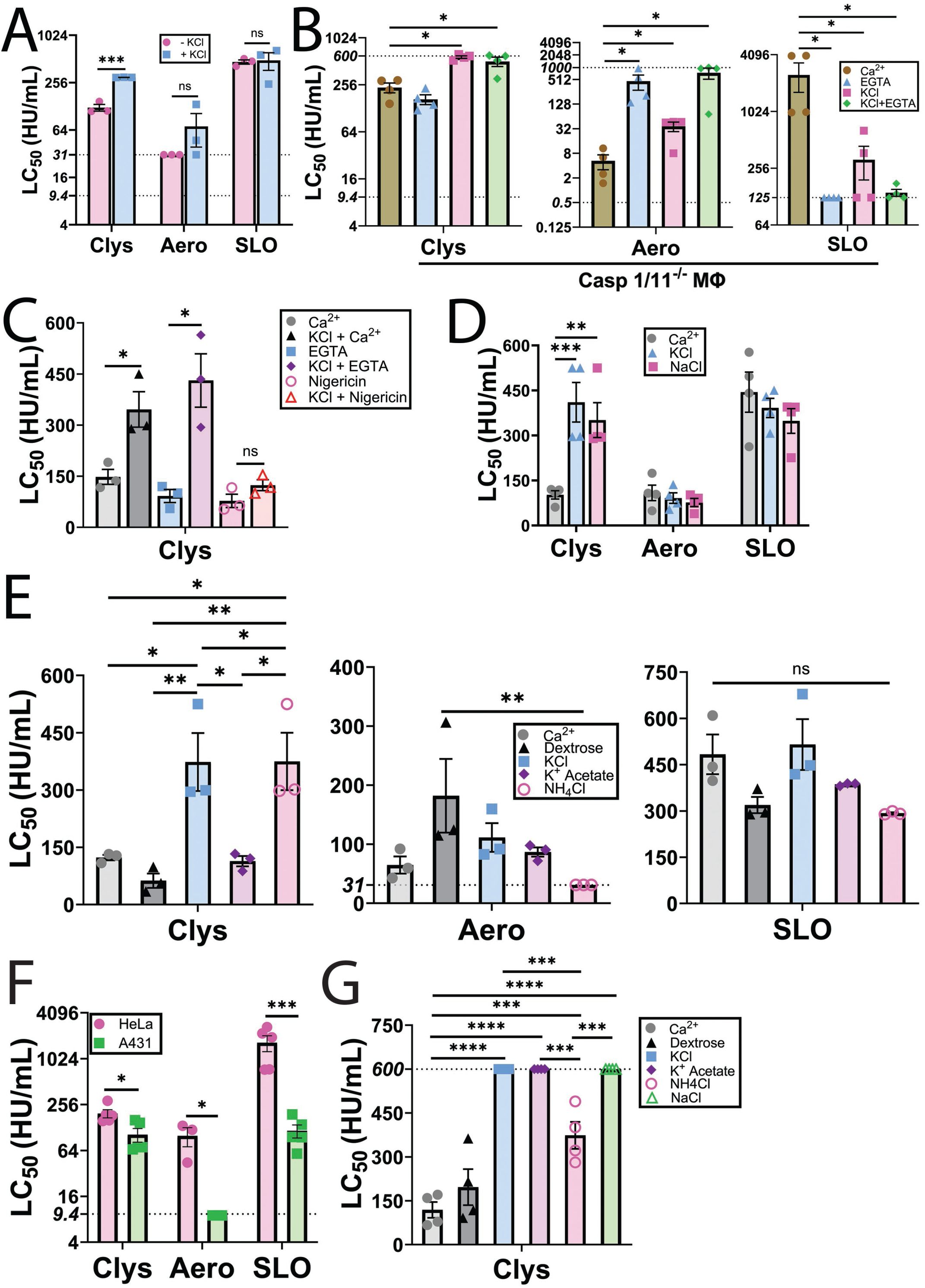
Chloride ions protect cells from candidalysin. (A) HeLa cells were challenged with 9.375-600 HU/mL candidalysin (Clys), 31-2000 HU/mL aerolysin (Aero), or 31-2000 HU/mL SLO in RPMI supplemented with or without 150 mM KCl. (B) Caspase 1/11^−/-^ bone-marrow derived macrophages (Casp1/11^−/-^ MΦ) were challenged with Clys, Aero, or SLO in RPMI supplemented with either 2 mM CaCl_2_ or 2 mM EGTA, and/or 150 mM KCl. (C) HeLa cells were challenged with Clys in RPMI supplemented with 2 mM CaCl_2_, 2 mM EGTA, 150 mM KCl and/or 20 μM nigericin alone or in combination. (D, E) HeLa cells were challenged with Clys, Aero, or SLO in RPMI supplemented with 2 mM CaCl_2_ and either: nothing, 150 mM NaCl, 150 mM KCl, 300 mM dextrose, 150 mM potassium acetate, or 150 mM NH_4_Cl. (F) HeLa or A431 vaginal epithelioid cells were challenged with 9.375-600 HU/mL Clys, 9.375-600 HU/mL Aero, or 31-2000 HU/mL SLO in RPMI supplemented with 2 mM CaCl_2._ (G) A431 cells were challenged with Clys in RPMI supplemented with 2 mM CaCl_2_ and either: nothing, 150 mM NaCl, 150 mM KCl, 300 mM dextrose, 150 mM potassium acetate, or 150 mM NH_4_Cl. PI uptake was analyzed by flow cytometry. The dotted lines indicates the upper (600 HU/mL for Clys, 1000 HU/mL for Aero) and lower limits (9.4 HU/mL for Clys, 125 HU/mL for SLO, 31 HU/mL, 9.4 HU/mL or 0.5 HU/mL for Aero) of detection. The LC_50_ was calculated as described in the methods. Graphs display independent experiments and mean ± S.E.M. of three (A, C, F: Aero), four (B, D, G), or five (F: Clys, SLO) five experiments. * p <0.05, ** p<0.01, *** p<0.001, **** p<0.0001, ns not significant by (A, B, D-G) repeated-measures one-way ANOVA with Tukey’s multiple comparison test or (C) repeated-measures two-way ANOVA with Geisser-Greenhouse correction and Šídák’s multiple comparisons test.

We next tested the ability of enforced K^+^ flux to increase cell death in response to candidalysin. We treated HeLa cells with potassium/proton ionophore nigericin, or EGTA, with and without 150 mM KCl prior to candidalysin challenge. Nigericin does not interfere with candidalysin activity^23^. Nigericin sensitized cells to candidalysin (Fig 3C, Supplementary Fig S5C). However, nigericin also overcame the inhibition provided by KCl, suggesting that its proton ionophore capabilities contribute to sensitizing cells to candidalysin (Fig 3C, Supplementary Fig S5C). In contrast, KCl overcame the ability of Ca^2+^ chelation to sensitize cells to candidalysin. We interpret these results to suggest that ion flux plays an important role in cell death due to candidalysin.

To distinguish between the contribution of the K^+^ and Cl^−^ ions, we compared the ability of multiple salts to protect cells against toxins. We first challenged HeLa cells with toxin in the presence or absence of 150 mM KCl or NaCl. While neither salt protected cells from aerolysin or SLO, both salts afforded cells similar protection against candidalysin (Fig 3D, Supplementary Fig S5D). We next tested 300 mM dextrose to control for osmolarity, and several potassium and chloride salts at 150 mM. Cell sensitivity to SLO was unaffected by all salts tested (Fig 3E, Supplementary Fig S5E). Cell sensitivity to aerolysin trended down upon dextrose treatment, but increased with NH_4_Cl (Fig 3E, Supplementary Fig S5E). In contrast, cells were protected against candidalysin in the presence of KCl or NH_4_Cl, but not potassium acetate (Fig 3E, Supplementary Fig S5E). We interpret these data to indicate that extracellular chloride may protect cells from candidalysin, but not aerolysin or SLO.

We next validated these findings in a cell line that better models the vaginal epithelium, A431 cells. We compared the sensitivity of HeLa cells and A431 cells to candidalysin, aerolysin and SLO (Fig 3F, Supplementary Fig S5F). A431 cells were at least 10x more sensitive to both aerolysin and SLO compared to HeLa cells, while they were only 2x more sensitive to candidalysin (Fig 3F, Supplementary Fig S5F). We next determined if Cl^−^ also protected A431 cells from candidalysin. We found that NaCl, KCl and potassium acetate all provided robust protection from candidalysin (LC_50_ >600), while NH_4_Cl provided 2-3x increased protection (Fig 3G, Supplementary Fig S5G). In contrast, controlling for osmolarity using dextrose did not protect cells (Fig 3G, Supplementary Fig S5G). Based on these data, we conclude that Cl^−^ also protects A431 cells from candidalysin.

Since Cl^−^ provides protection in multiple cell lines, it is possible that Cl^−^ interacts with the positively charged residues in candidalysin to prevent binding and/or oligomerization. We tested this hypothesis by measuring the ability of candidalysin to perforate a supported lipid bilayer in the presence of each ion. Artificial lipid bilayers were created using diphytanoyl phosphatidylcholine (DPhPC) and the Orbit e16, and a voltage of -50 mV applied in order to mimic a host cell membrane. Changes in the current across the bilayers was recorded to monitor their stability. The addition of candidalysin induced changes to the current consistent with candidalysin insertion into the bilayer and rupture events, with no significant differences in latency or frequency observed between treatment conditions (Fig 4). No events occurred when bilayers were exposed to buffer in the absence of candidalysin (not shown). Based on these results, we conclude that Cl^−^ exerts its protective effects by targeting the host cell and not by directly interacting with candidalysin to prevent activity.

**Figure 4.**
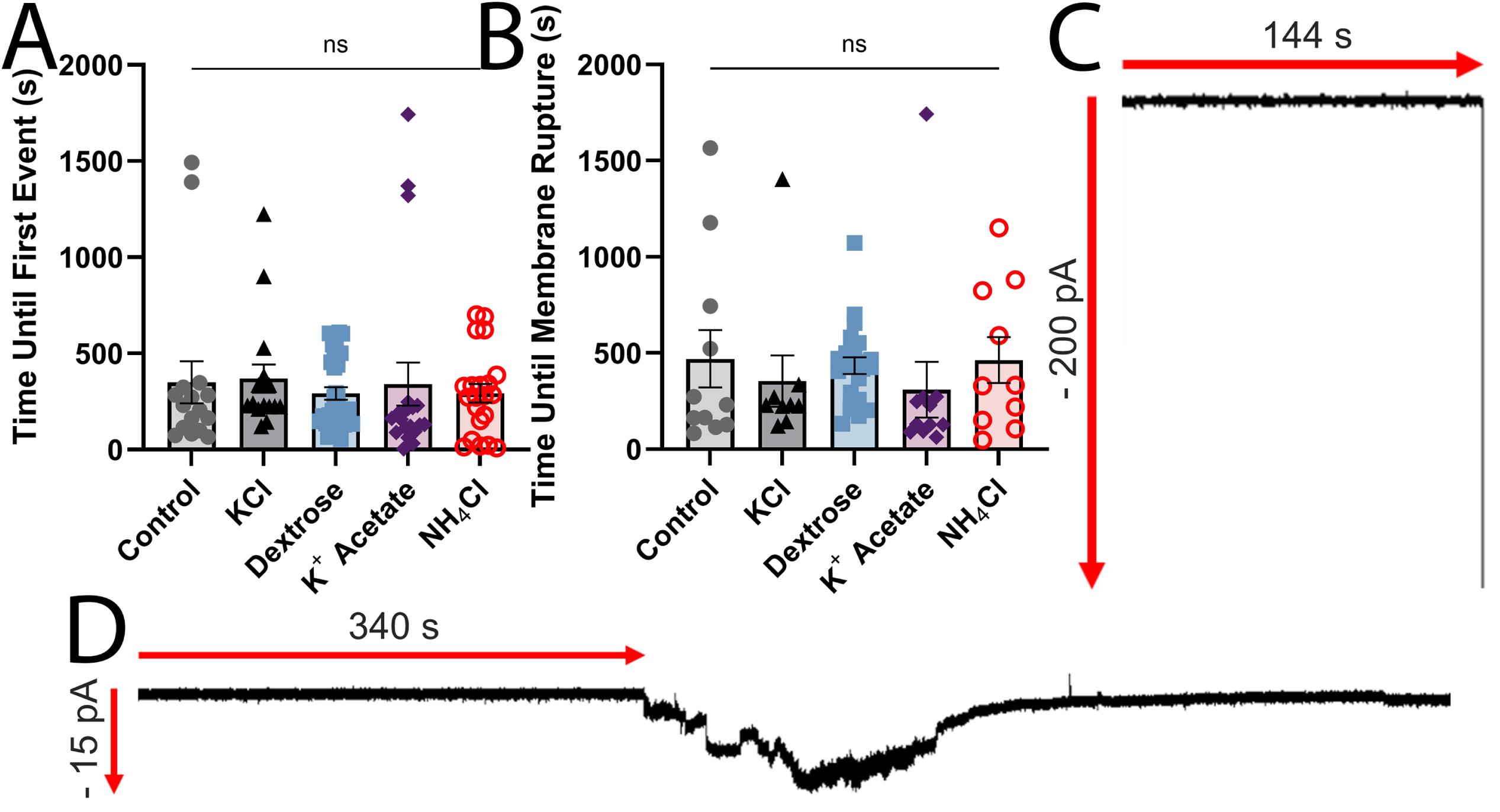
Candidalysin displays consistent artificial membrane interactions. Artificial lipid bilayers were exposed to 10 μM candidalysin in buffers containing 110 mM chloride ions and 5 mM potassium ions (control), and either: 150 mM KCl, 300 mM dextrose, 150 mM potassium acetate, or 150 mM NH_4_Cl. Time upon candidalysin exposure to (A) first event, defined as either candidalysin insertion into the bilayer or bilayer rupture, and (B) bilayer rupture. Graphs show independent data points, and the mean ± S.E.M. One representative trace of rupture (C) and insertion (D) is presented, from N = 16 (control), N = 16 (KCl), N = 31 (dextrose), N = 20 (potassium acetate), N = 21 (NH_4_Cl). Data were statistically analyzed by ANOVA with Šidák post-hoc for multiple comparisons.

### Candidalysin triggers MEK dependent repair

We next tested the role of MEK signaling in candidalysin resistance. MEK is activated by candidalysin, and signaling via ERK is important for cell survival at late time points^23^, so it is possible that the ERK-independent signaling protects cells at early time points like it does for CDCs^17^. We blocked MEK activation by pretreating with the MEK inhibitor U0126 for 30 min (Fig 5A), which is specific for MEK in this system^17^. MEK inhibition increased the sensitivity of HeLa cells to both candidalysin and SLO, but not to aerolysin (Fig 5B-D, Supplementary Fig S6). When compared to the total Ca^2+^-dependent repair, MEK inhibition accounted for ∼40% of Ca^2+^-dependent repair against candidalysin and ∼50% of Ca^2+^-dependent repair against SLO. Overall, candidalysin triggers MEK-dependent repair, though to a lesser extent than CDCs.

**Figure 5.**
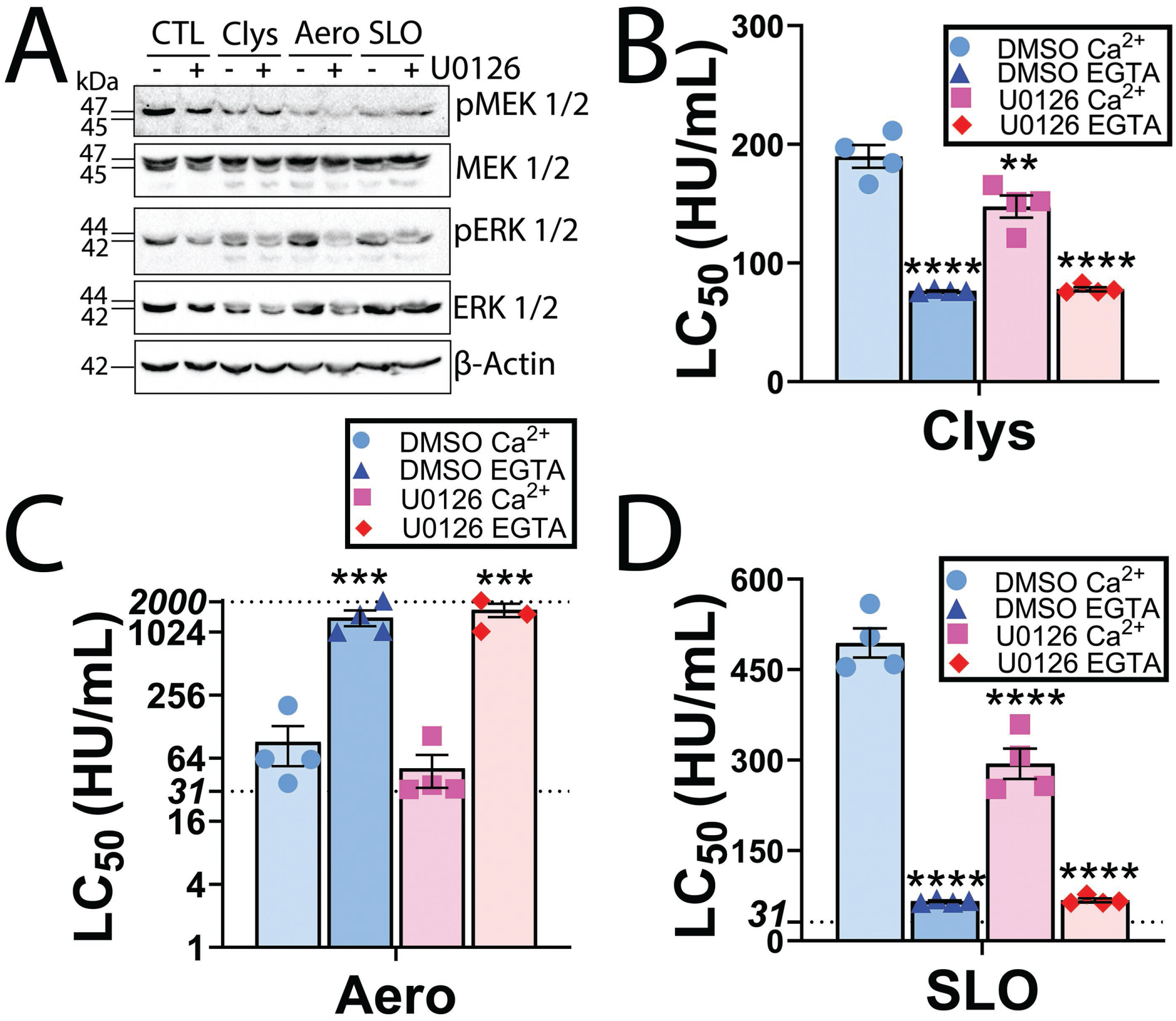
MEK activation provides limited protection against candidalysin. HeLa cells were serum starved for 30 min and pre-treated with 20 μM MEK inhibitor U0126 or DMSO for 30 min. (A) Cells were then challenged with sublytic toxin concentrations (50 HU/mL candidalysin (Clys), 62 HU/mL aerolysin (Aero) or 250 HU/mL SLO)) in the continued presence of U0126 or DMSO and analyzed by western blot. Blots were probed with the indicated antibodies and HRP-conjugated secondary antibodies. (B-D) Serum starved and DMSO or U0126-treated Hela cells were challenged with (B) 9.375-600 HU/mL Clys, (C) 31-2000 HU/mL Aero, or (D) 31-2000 HU/mL SLO in RPMI supplemented with either 2 mM CaCl_2_ or 2 mM EGTA. PI uptake was analyzed by flow cytometry. The LC_50_ was calculated as described in the methods. The dotted lines indicate the upper and lower limits of detection. One representative blot from three independent experiments is shown. Graphs show four independent experiments, and the mean ± S.E.M. ** p<0.01, *** p<0.001, **** p<0.0001 by repeated-measures one-way ANOVA with Tukey’s multiple comparison test compared to DMSO Ca^2+^.

### Annexins provide limited and delayed protection against candidalysin

We further explored the repair pathway downstream of MEK activation. One downstream target of MEK-dependent repair is annexin A2 (A2) ^17^. Annexins A1 and A6 are independent of MEK activation, but are involved in membrane repair^14, 17, 37^. We tested the role of these three annexins in the repair of candidalysin-mediated damage using siRNA. Annexin knockdown efficiency was 70% for A1, 85% for A2, and 60% for A6 (Fig. 6A-B). Knockdown of each annexin sensitized cells to candidalysin, aerolysin and SLO (Fig 6C, Supplementary Fig S7).

**Figure 6.**
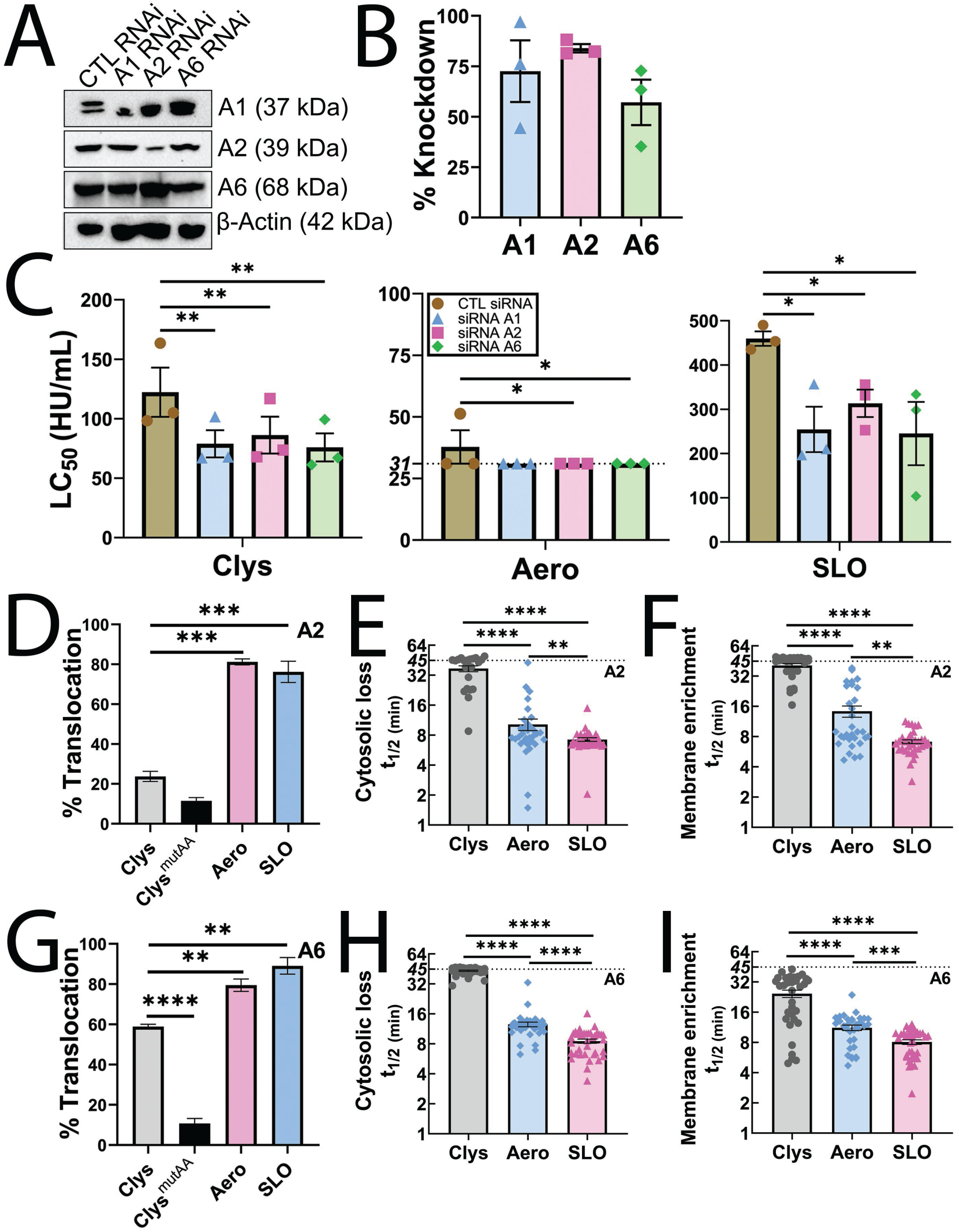
Annexins provide limited protection against candidalysin. (A-C) HeLa cells were transfected with control (CTL), annexin A1 (A1), annexin A2 (A2) or annexin A6 (A6) siRNA for 3 days and then (A, B) lysed for western blot analysis, or (C) challenged with 9.375-600 HU/mL candidalysin (Clys), 31-2000 HU/mL aerolysin (Aero), or 31-2000 HU/mL SLO. PI uptake was analyzed by flow cytometry. The LC_50_ was calculated as described in the methods. (A) Portions of the blot were probed with antibodies against A1, A2, A6 or β-actin followed by HRP-conjugated secondary antibodies. (B) Blots were quantified by densitometry and the knockdown efficiency compared to control siRNA determined. (D-F) A2-GFP or (G-I) A6-YFP transfected HeLa cells were challenged with sublytic toxin concentrations (50 HU/mL Clys, 62 HU/mL Aero, or 250 HU/mL SLO) or a mass equivalent of candidalysin^AA^ (Clys^mutAA^). Cells were imaged by confocal microscopy for ∼45 min at 37°C and then lysed using 1% Triton-X-100. The (D, G) fraction of cells showing annexin membrane translocation is shown. For cells showing annexin translocation, the time to half maximal (t_1/2_) (E, H) cytosolic depletion or (F, I) membrane accumulation of A2 or A6 is shown. The dotted lines indicate the upper and lower limits of detection. The blots show one representative experiment from three independent experiments. Graphs show the mean ± S.E.M. Individual data points represent (B) each of three independent experiments or (E, F, H, I) individual cells from 3 independent experiments. * p <0.05, ** p<0.01, *** p<0.00, **** p<0.0001 by (C, D, G) repeated measures or (E, F, H, I) regular one-way ANOVA with Tukey’s multiple comparison test.

These results suggest that annexins promote cellular resistance to candidalysin. We further explored the contribution of annexins to repair against candidalysin using high-resolution live cell imaging. A2 translocates rapidly to the plasma membrane^20^, except when MEK is inhibited^17^. Annexin membrane recruitment is faster for CDCs compared to aerolysin^21^. We challenged A2-GFP or A6-YFP transfected cells with candidalysin, candidalysin^AA^ as a control, aerolysin or SLO, and imaged annexin dynamics (Fig 6D-I, Supplementary Fig S8A, B, Videos 1 & 2). Annexins translocated to the membrane in ∼20% of A2-GFP+ cells and ∼60% A6-YFP+ cells after candidalysin challenge, compared to translocation in 75-90% of cells challenged with aerolysin or SLO challenge (Fig 6D, G). To measure cell viability during annexin translocation, we analyzed TO-PRO3 uptake. Annexins translocated to the membrane without TO-PRO3 accumulation (Supplementary Fig S8C, Videos 1 & 2). We measured annexin translocation by determining the half time (t_1/2_) of depletion of the annexin from the cytosol, plus the half time (t_1/2_) of annexin enrichment on the plasma membrane. Similar to previous results^17, 21^, the t_1/2_ for A2 and A6 depletion and recruitment was ∼7 min and ∼8 min, respectively, for SLO, and ∼12 min for aerolysin (Fig 6E, F, H, I, Supplementary Fig S8A, B, Videos 1 & 2). In contrast, active candidalysin triggered delayed cytosolic depletion and membrane recruitment of both A2 and A6, with a t_1/2_ of ∼40 min cytoplasmic annexin depletion and membrane recruitment of A2 (Fig 6E, F, H, I, Supplementary Fig S8A, B, Videos 1 & 2). Surprisingly, A6 was recruited to the membrane faster (t_1/2_ ∼25 min) than A2 during candidalysin challenge (Fig 6E, F, H, I, Supplementary Fig S8A, B, Videos 1 & 2). This is similar to the situation when MEK is not active in repair^17^. Taken together, we interpret these data to suggest that annexins contribute to candidalysin resistance, but they are recruited later in the repair process than for other toxins.

### Microvesicle shedding removes candidalysin from cells

Since annexin clogging and patch repair had minimal to no impact on repair, we tested the hypothesis that candidalysin is shed on microvesicles. We challenged cells with a sublytic dose of each toxin, or candidalysin^AA^ to control for spontaneous vesicle release. All three active toxins were shed on microvesicles, though oligomeric aerolysin also remained present on the cell surface (Fig 7A). Since candidalysin^AA^ was not detected in the blot, we determined if the antibody could recognize candidalysin^AA^. We performed dot blots using either wild-type candidalysin, candidalysin^AA^, or aerolysin. The anti-candidalysin antibody recognized both candidalysins, but not aerolysin, while the anti-aerolysin antibody only recognized aerolysin (Fig 7B).

**Figure 7.**
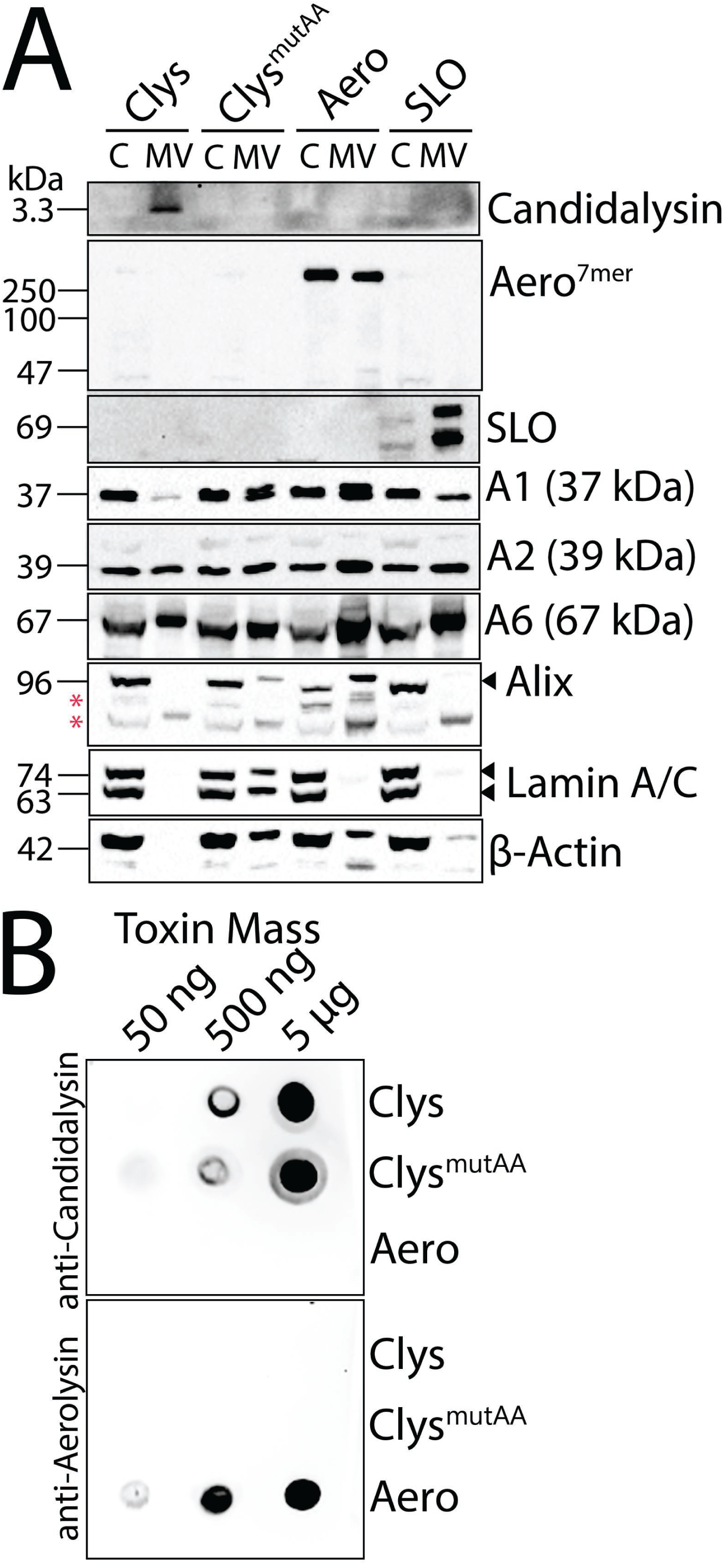
Candidalysin is removed via membrane shedding. (A) HeLa cells were challenged with sublytic toxin concentrations (50 HU/mL candidalysin (Clys), 62 HU/mL aerolysin (Aero) or 250 HU/mL SLO) or a mass equivalent of candidalysin^AA^ (Clys^mutAA^) for 1 h at 37°C. Cells were centrifuged at 2000xg for 5 min to yield cell pellets (denoted C). Cell supernatants were spun at 100,000xg for 40 min at 4°C for the microvesicle pellet (MV). Proteins were resolved by SDS-PAGE and analyzed by western blot using the indicated primary antibodies. The blots show one representative experiment from four independent experiments. (B) Clys, Clys^mutAA^, or Aero were spotted on nitrocellulose and analyzed by western blot using anti-candidalysin or anti-aerolysin antibodies. One representative blot of three independent experiments is shown. * indicates non-specific bands.

We then determined if other cellular proteins are shed. Along with the active toxins, A2 and A6 were shed in microvesicles for all three toxins (Fig 7A), consistent with the previous results^19, 21^. Neither A1 nor the Endosomal Sorting Complex Required for Transport (ESCRT) protein ALG-2-interacting protein X (ALIX) were shed following candidalysin challenge, though they were shed in response to aerolysin (Fig 7A). To distinguish between shed vesicles and cellular debris from lysed cells, we probed for the nuclear proteins Lamin A/C. Lamin A/C was not shed from cells challenged with active toxins (Fig. 7A), indicating that the microvesicle fraction did not contain detectable cellular debris. Thus, candidalysin is shed on microvesicles by cells.

## Discussion

In this study, we determined which repair mechanisms respond to candidalysin and compared these responses to pore-forming toxins from the CDC and aerolysin families. We found that candidalysin triggered a distinct repair profile compared to both CDCs and aerolysin. Similar to CDCs, candidalysin triggered Ca^2+^-dependent repair and microvesicle shedding. However, candidalysin triggered less MEK-dependent repair than CDCs. In contrast to both aerolysin and CDCs, extracellular chloride, but not patch repair, protected cells from candidalysin. Finally, annexin recruitment was reduced and delayed compared to other toxins. Based on our data, we conclude that candidalysin triggers distinct molecular mechanisms to drive microvesicle shedding, which suggests approaches to bolster specific membrane repair mechanisms could reduce *Candida* pathogenicity.

Our present work integrates and expands the repair mechanisms used to resist candidalysin. Our results are consistent with prior work^3, 22, 23^ showing the Ca^2+^-dependence of repair resisting candidalysin. We find additional active repair mechanisms that resist candidalysin beyond ESCRT-III mediated shedding^22^ and autophagy^15^, including a chloride trigger, and involvement of MEK and annexins. Since MEK inhibition had partial effect on repair, and annexin translocation was delayed, shedding via ESCRT activation and via MEK might represent two distinct repair pathways that both trigger microvesicle shedding. Interestingly, prior work observed Alix and Galectin-3 recruitment to roughly half of the sites of *Candida albicans* penetration^15^, suggesting that the other sites were either undamaged, or that different repair pathways were engaged. Where ESCRT pathways diverge from other repair pathways downstream of Ca^2+^ influx remains to be determined.

Our findings here differ from others’ results in the role of lysosomal exocytosis and/or patch repair to resist candidalysin. Prior work observed recruitment of lysosomal proteins to the site of damage^15, 22^, while our dysferlin assay showed no contribution of dysferlin to protection. Since the prior work used *Candida albicans* for damage instead of candidalysin alone, another protein or factor could trigger patch repair in those systems. Another possibility is that dysferlin recruitment/involvement fails to capture the full spectrum of patch repair, so assay-specific differences could account for the different results. Finally, it is possible that lysosomal exocytosis contributes to the cellular return to homeostasis, instead of directly contributing to repair of candidalysin-induced lesions. Our assay is focused on repair because it measures cell viability instead of protein relocalization, so it would not capture post-repair membrane rearrangements. In summary, our findings provide a novel perspective to cellular repair responses to candidalysin

One key novel finding was that increasing the extracellular Cl^−^ concentration protected cells from candidalysin, in contrast to any role of Cl^−^ for SLO or aerolysin. Several lines of evidence support Cl^−^ as the active ion over K^+^ or changes in osmolarity, despite the role of K^+^ channels in cell death^38^. First, prior work using glibenclamide to block K^+^ channels did not impact damage due to candidalysin^39^. Second, addition of dextrose or potassium acetate failed to enhance cellular resistance to candidalysin. Third, nigericin-mediated death in our assays was not blocked by extracellular K^+^. Finally, multiple Cl^−^ salts provided protection, while salts without Cl^−^ and dextrose all failed to protect cells. While it remains possible that monovalent cations collectively provide protection, we note that the electrochemical gradients driven by Na^+^ and K^+^ run in opposite directions^38^, yet both NaCl and KCl provided similar protection. In contrast, the typical extracellular Cl^−^ concentration is ∼2-3 fold higher than intracellular concentrations^40^. How increasing extracellular Cl^−^ improves cellular resistance remains to be determined.

Our findings place renewed interest in tracking Cl^−^ in tissues. The ventricles of the heart maintain 200-340 mM Cl^−41^. Systemically, Cl^−^ varies in the vaginal mucosa with the menstrual cycle, in the oral mucosa with diet and hydration, and is elevated in response to bicarbonate ion loss in the gut caused by diarrhea^41, 42^. Interestingly, vaginal Cl^−^ may be reduced during mycosis^42^, which could be a pathogen response. While any specific mechanisms blocking or enhancing Cl^−^ secretion during *C. albicans* infection remain to be determined, the Cl^−^ concentrations we used *in vitro* may be physiologically relevant in localized areas during *C. albicans* infection.

There are some potential mechanisms by which Cl^−^ could enhance resistance to candidalysin. Chloride salts like ammonium chloride can promote autophagy^43^. Given the role for autophagy in repairing candidalysin lesions^15^, it is possible that extra Cl^−^ enhances autophagy, which drives repair. Alternatively, chloride channels could promote repair. Proteins in the anoctamin family (TMEM16A-F) include Ca^2+^-dependent Cl^−^ channels^44–46^. For example, Anoctamin 6/TMEM16F contributes to the outwardly rectifying chloride channel and is involved in cell death and size^46^. Anoctamin 6 is previously implicated in promoting repair against both listeriolysin O and SLO^45^. The counterargument is that we observed an effect for Cl^−^ against candidalysin, but not SLO. Other species of *Candida* produce candidalysins that are more potent than *C. albicans*^4^. While it is likely that these candidalysins are repaired by similar mechanisms, it remains to be determined if different levels of Cl^−^ are needed for protection. Thus, the mechanism by which increased extracellular Cl^−^ enhances repair against candidalysin is a future avenue for research.

Another novel finding was that annexins had a delayed response to candidalysin compared to SLO or aerolysin. Taken with our findings that knockdown of either A1, A2 or A6 contributed mild effects to reducing repair, we propose that annexins serve as a backup mechanism to resist candidalysin. In many cases, A6 translocated faster than A2, which further suggests that MEK dependent repair is not the primary means of resisting candidalysin. Alternatively, candidalysin could trigger a programmed cell death pathway, and annexin translocation is either dampened by the death pathway, or activation delayed until just prior to cell lysis. Future directions include determining the interplay between programmed cell death pathways and membrane repair.

Our work had several limitations due to the experimental system. We did not test candidalysin in the context of the whole pathogen, or in *in vivo* infection models. We used multiple cell lines and primary cells, which have physiological differences. We used erythrocytes and HeLa cells to standardize cellular responses because these are well-characterized systems in which to measure toxin-dependent repair mechanisms. We used myoblasts to test the contribution of dysferlin knockout because dysferlin is not expressed in HeLa cells. Finally, we used primary macrophages and vaginal epithelioid cells because these are the cell types targeted by *C. albicans*. In some cases like macrophages, the resistance to certain toxins is different than HeLa cells. We relied on prior work^15, 22^ for the contribution of autophagy and ESCRT to repair against candidalysin. While we found an effect from Cl^−^ to enhance repair, the signaling intermediates between Cl^−^ and repair remain a future research direction. Similarly, the toxin determinants that determine which repair pathways are activated remains unknown.

In summary, we compared repair responses between candidalysin and toxins from two other families, CDCs and aerolysins. Candidalysin triggered a distinct repair profile compared to these toxins that included Ca^2+^-dependent repair, microvesicle shedding and resistance in the presence of elevated chloride levels. In contrast to other toxins, annexin recruitment was reduced and delayed, dysferlin-mediated patch repair did not contribute to repair, and MEK-dependent repair was reduced. Overall, we conclude that candidalysin triggers distinct membrane repair mechanisms that could be enhanced to reduce *C. albicans* pathogenicity.

## Materials and Methods

### Reagents

All reagents were from Thermo Fisher Scientific (Waltham, MA, USA) unless otherwise noted. The MEK1/2 inhibitor U0126 was from Tocris (Minneapolis, MN, USA) (Cat# 1144) or Alfa Aesar (Cat# J61246). Negative control siRNA or Silencer pre-designed siRNAs for annexin A1 (Assay ID: 146988), A2 (Assay ID: 147285) or A6 (Assay ID: 241008) were from Invitrogen (Waltham, MA, USA). Propidium iodide (PI) was from Biotium (Fremont, CA, USA) (Cat # 40016). MTT reagent (Cat# 2809-1G) was from BioVision (Milpitas, CA, USA). D-threo-1-phenyl-2-hexadecanoylamino-3-morpholino-1-propanol ·HCl (PPMP) was from Avanti Polar Lipids (Alabaster, AL, USA) (catalog no. 870792).

### Antibodies

The anti-annexin A1 monoclonal antibody (mAb) clone CPTC-ANXA1-3-s was deposited to the Developmental Studies Hybridoma Bank (DSHB) by Clinical Proteomics Technologies for Cancer (DSHB Hybridoma Product CPTC-ANXA1-3) and anti-Lamin A/C mAb clone MANLAC-4A7-s was deposited to the DSHB by GE Morris (DSHB Hybridoma Product MANLAC1(4A7)). Both were obtained from the DSHB, created by the NICHD of the NIH and maintained at the University of Iowa, Department of Biology (Iowa City, IA, USA). Anti-β-actin mAb clone AC-15 (Cat# GTX26276) was from GeneTex (Irvine, CA, USA). Anti-SLO mAb clone 6D11 (catalog: NBP1-05126) was from Novus Biologicals (Littleton, CO, USA). Anti-Alix mAb clone 3A9 mAb was from BioLegend (San Diego, CA, USA). The anti-aerolysin rabbit polyclonal antibody^21^ and myc-tagged CAL1-H1 anti-candidalysin camelid nanobody^47^ were previously described. Anti-MEK (Cat# 9122L), anti-phospho-MEK [Ser217/Ser221] (Cat# 9121S), anti-ERK p44/42 (Cat# 9102S), and anti-phospho-ERK [Thr202/Tyr404] (Cat# 9101S) rabbit polyclonal antibodies were from Cell Signaling Technologies (Danvers, MA, USA). Anti-mouse-IgG (Cat# 711-035-151), anti-rabbit-IgG (Cat# 711-035-152), or anti-alpaca IgG (Cat# 128-035-230) conjugated to horse radish peroxidase (HRP) were from Jackson ImmunoResearch (West Grove, PA, USA).

### Plasmids

The pET22b plasmids encoding His-tagged aerolysin and aerolysin*^Y^*^221^*^G^* were kind gifts from Gisou van der Goot (École Polytechnique Fédérale de Lausanne, Canton of Vaud, Switzerland) ^48^. The wild-type aerolysin contains Q254E, R260A, R449A and E450Q mutations and the C-terminal KSASA was replaced with NVSLSVTPAANQLE HHHHHH compared to sequences in GenBank (i.e. M16495.1). Human annexin A6 fused to YFP was a generous gift from Annette Draeger (University of Bern, Bern, Switzerland) ^49^. Human annexin A2-GFP (Addgene plasmid no. 107196) was from Addgene^50^. OlyA fused to mCherry in pET21c(+) was a kind gift from Kristina Sepčić (University of Ljubljana, Ljubljana, Slovenia) ^51^, and the E69A mutation was introduced as described^17^. Cysteine-less, His-tagged wild-type and G398V/G399V monomer-locked (ML) SLO codon-optimized for *Escherichia coli* expression^17^ and GFP-dysferlin^21^ were previously described.

### Mice

All experimental mice were housed and maintained according to Texas Tech University Institutional Animal Care and Use Committee (TTU IACUC) standards, adhering to the Guide for the Care and Use of Laboratory Animals (8th edition, NRC 2011). The TTU IACUC approved mouse use. Casp1/11^−/-^ mice on the C57BL/6 background were purchased from the Jackson Laboratory (Bar Harbor, ME, USA) (stock #016621). Mice of both sexes aged 6–15 weeks were used to prepare bone-marrow derived macrophages (BMDM). Sample size was determined as the minimum number of mice needed to provide enough bone marrow for experiments. Consequently, no randomization or blinding was needed. Mice were euthanized by asphyxiation through the controlled flow of pure CO_2_ followed by cervical dislocation.

### Cell culture

All cell lines were maintained at 37°C, 5% CO_2_. HeLa cells (ATCC (Manassas, VA, USA) CCL-2) and A431 cells (ATCC CRL-1555) were cultured in DMEM (Corning, Corning, NY, USA) supplemented with 10% Equafetal serum blend (Atlas Biologicals, Fort Collins, CO, USA) and 1x L-glutamine (D10 medium). Casp1/11^−/-^ BMDM were isolated from bone marrow and differentiated for 7-21 days in DMEM supplemented with 30% L929 cell supernatants, 20% fetal calf serum (VWR Seradigm, Radnor, PA, USA), 1 mM sodium pyruvate and 1x L-glutamine, as previously described^17, 19^.

### Toxins

Wild-type (SIIGIIMGILGNIPQVIQIIMSIVKAFKGNK) candidalysin and candidalysin^AA^ (SIIGIIMGILGNIPQVIQIIMSIVKAF**A**GN**A**) were synthesized by Peptide Protein Research, Ltd (Hampshire, UK) as previously described^3^. SLO, OlyA-mCherry, and pro-aerolysin were induced in *E. coli* BL21 GOLD cells and purified as previously described^17, 21, 52^. Briefly, log phase cells were induced with 0.2% arabinose (SLO, SLO ML), or 0.2 mM IPTG (OlyA-mCherry, OlyA*^E69A^*-mCherry, pro-aerolysin, and pro-aerolysin*^Y^*^221^*^G^*) for 3 h at room temperature, frozen, lysed, and then purified using Nickel-NTA resin. Protein concentration was determined by Bradford Assay (Supplementary Table S1). Purity was assessed by SDS-PAGE. The hemolytic activity of each toxin was determined as previously described^17, 21^ using human red blood cells (Zen-Bio, Research Triangle Park, NC, USA) (Supplementary Table S1). One hemolytic unit (HU) is defined as the quantity of toxin required to lyse 50% of a 2% human red blood cell solution in 30 min at 37°C in 2 mM CaCl_2_, 10 mM HEPES, pH 7.4 and 0.3% BSA in PBS^17, 21, 52^. We used HU/mL to normalize toxin activities across experiments and to achieve consistent cytotoxicity across toxin preparations. Sublytic doses were defined as the highest toxin concentration that killed <20% of target cells. For HeLa cells, the sublytic dose was 250 HU/mL for SLO, 62 HU/mL for aerolysin and 50 HU/mL for candidalysin. Pro-aerolysin was activated using trypsin as previously described^21^ prior to assays.

### Transfection

HeLa cells were plated at 2 × 10^5^ cells per 35 mm glass-bottom dish or per well in a 6 well plate and transfected with 750 ng annexin A6, annexin A2, peGFP-N1 or GFP-Dysferlin using Lipofectamine 2000 in Opti-MEM two days before imaging or cytotoxicity assays. D10 media was replaced on the day after transfection. Transfection efficiencies for each construct ranged between 65-80%. For RNAi, HeLa cells were plated at 2 × 10^5^ cells in 6-well plates and transfected the following day with 10 nM siRNA using Lipofectamine 2000 in Opti-MEM for 48-72 h as previously described^17, 21^.

### MTT/LDH Assays

The MTT [3-(4,5-dimethylthiazol-2-yl)-2,5-diphenyltetrazolium bromide] and LDH (lactate dehydrogenase) assays were performed and analyzed as previously described^21^. Briefly, HeLa cells were harvested and resuspended at 1 × 10^5^ cells per well in a 96 well flat bottom plate in DMEM (without Phenol Red) supplemented with 2 mM CaCl_2_. Toxins were diluted in DMEM according to hemolytic activity (wild-type toxins) or equivalent mass (non-hemolytic toxins), serially diluted two-fold, added to wells containing cells, and incubated for 30 min, 1 h or 2 h at 37°C. Triton-X-100 was added to a final concentration of 0.5% to four wells containing cells as a positive control for maximum LDH release and incubated for 30 min. After incubation, cells were centrifuged at 1200× g for 5 min at 4°C to pellet cells. Cell pellets were assayed for MTT activity, while supernatants were assayed for LDH activity. For the MTT assay, cells were resuspended in 1.1 mM MTT assay reagent in DMEM without Phenol Red and incubated at 37°C for 4 h. Formazan was dissolved in SDS-HCl at 37°C overnight. A_570_ was measured with a plate reader (Bio-Tek, Winooski, VT, USA). The % viable cells was calculated as follows: % Viable = (Sample − Background)/(Control − Background) × 100. Then, % Specific Lysis was calculated as 100% − Viable. For the LDH assay, cell supernatants were assayed per the manufacturer’s instructions. % LDH release was calculated as (Sample − Background)/ (Maximum LDH release − Background) × 100.

### Flow Cytometry Assay

Cytotoxicity was performed as described^17, 21, 52^. Briefly, cells were transfected for 48-72 h or incubated with 40 µM PPMP for 72 h to promote ceramide accumulation^53^. 1×10^5^ cells were challenged in suspension with various toxin concentrations for 30 or 60 min at 37°C in RPMI supplemented with 2 mM CaCl_2_ and 20 μg/mL PI. Cells were analyzed on a 4-laser Attune Nxt flow cytometer. Assay variations included changes to the media (replacing 2 mM CaCl_2_ with 2 mM EGTA, adding 150 mM KCl, 150 mM NaCl, 300 mM Dextrose, 150 mM potassium acetate, 150 mM NH_4_Cl, or 20 μM nigericin), or serum-starving cells in DMEM for 30 min at 37° C and pretreating with DMSO or 20 µM U0126 in serum-free media for 30 min at 37° C. For analysis of cell lysis, we gated out the debris and then quantified the percentage of cells with high PI fluorescence (2–3 log shift) (PI high), or background fluorescence (PI negative). We calculated specific lysis as: % Specific Lysis = (% PI High^Experimental^ − % PI High^Control^)/ (100 − % PI High^Control^) × 100. The toxin dose needed to kill 50% of cells was defined as the Lethal Concentration 50% (LC_50_) and was determined by logistic modeling using Excel (Microsoft, Redmond, WA, USA) as previously described^26^.

### Fluorescent annexin shedding assays

Annexin translocation assays were performed as previously described^17, 21^. Briefly, HeLa cells were plated at 2 × 10^5^ cells per 35 mm glass-bottom dish and transfected with 750 ng A6-YFP or A2-GFP using Lipofectamine 2000 two days prior to imaging. The transfection efficiency was ∼65-80%. Transfected cells were challenged with a sublytic toxin dose for active toxins (250 HU/mL SLO, 62.5 HU/mL aerolysin or 50 HU/mL candidalysin) or mass equivalent of candidalysin^AA^ (candidalysin^mutAA^) and imaged at 37°C in RPMI, 25 mM HEPES pH 7.4, and 2 mM CaCl_2_ with 2 μg/mL TO-PRO3 for 45 min using a Fluoview 3000 confocal microscope (Olympus, Tokyo, Japan) equipped with a 60×, 1.42 NA oil immersion objective and recorded using the resonant scanner at 1–3 s per frame. A6-YFP and A2-GFP were excited using a 488 nm laser, while TO-PRO3 was excited with a 640 nm laser.

After 45 min, the assay was terminated by adding an equal volume of 2% Triton-X-100 to give a final concentration of 1%. The percentage of cells showing fluorescent annexin translocation was determined by comparing the intensity at 15 min to the intensity 1 min after toxin addition. If these values were below 80% of the initial value, cells were considered to show annexin translocation. In cells showing fluorescent annexin translocation, the extent of fluorescent annexin depletion from cells was assessed by measuring the intensity from the middle of the cell over time by plotting Z profiles in FIJI (NIH, Bethesda, MD, USA). Membrane accumulation of annexins (A2 or A6) over time was calculated by determining the changes in the fluorescence intensity along the edges of the cells of translocated cells by plotting the Z profile in FIJI. The intensity was then normalized on a per-cell basis and individually analyzed cells were plotted.

TO-PRO3 uptake was determined in fluorescent annexin translocated cells by plotting the Z profile of a nuclear subsample over time. The data were normalized to the integrated intensity of the brightest point. The t_1/2_ for annexin cytosolic depletion, membrane accumulation, or TO-PRO3 uptake was calculated by measuring the half-maximal intensity between the starting intensity and the average intensity of the last 4 min prior to Triton-X-100 addition. From the multi-plane images, >25 cells were analyzed from at least three independent experiments and graphed using Microsoft Excel. Quantified values for individual cells were represented with Prism 10.4.1 (GraphPad, San Diego, CA, USA).

### Bleach correction and supplemental videos

Images were exported as .tif and split into individual monochromatic red, green, and blue channels in FIJI. For display (but not analysis), bleach correction was performed as previously described^17^ on the “green” channel cells by histogram matching using FIJI followed by a median pass filter. The monochromatic images were then merged to form RGB .tif files, time-stamped, annotated, and exported as AVIs for supplementary videos.

### Bilayer permeabilisation assay

Artificial lipid bilayer permeabilisation experiments were performed using the Orbit e16 (Nanion) and Elements Data Reader (3.8.7) software. To produce bilayers, DPhPC lipids were dissolved in octane to 25 mg/mL, then applied to multielectrode cavity array (MECA) chips featuring 16 microelectrode cavities (Ionera). The bilayer was coated in solution containing 110 mM chloride ions and 5 mM potassium ions, and either: nothing, 150 mM KCl, 300 mM dextrose, 150 mM potassium acetate, or 150 mM NH_4_Cl. A voltage of -50 mV was applied and candidalysin was added to the well to a final concentration of 10 μM. Current (pA) was recorded for 30 min at room temperature. Data were analyzed using Clampfit software v10.3 (Molecular Devices).

### Ultracentrifugation

Ultracentrifugation was performed as previously described^17^. Briefly, 5 ×10^6^ - 10 ×10^6^ cells were challenged with a sublytic dose (50 HU/mL for candidalysin, 62 HU/mL for aerolysin, 250 HU/mL for SLO) for 1 h at 37° C, and collected by centrifugation at 2000xrpm for 5 min (cell pellet). The supernatant was centrifuged at 100,000xg for 40 min at 4° C. The high-speed pellet was collected as the microvesicle fraction. Cells and microvesicles were solubilized in 95° C 1x SDS-sample buffer, heated for 5 min, and cells sonicated to shear DNA.

### SDS-PAGE and immunoblotting

Cells or proteins were dissolved in 95° C 1x SDS-sample buffer, heated for 5 min at 95° C and sonicated. Samples were resolved on 10% polyacrylamide gels at 170 V for ∼90 min and transferred to nitrocellulose in an ice bath with transfer buffer (15.6 mM Tris and 120 mM glycine) at 110 V for 90 min. Blots were blocked overnight in 5% skim milk in 10 mM Tris-HCl, 150 mM NaCl and 0.1% Tween-20 at pH 7.5 (TBST). For dot blots, 5 µg, 0.5 µg, or 50 ng candidalysin, candidalysin^AA^ or aerolysin were spotted on nitrocellulose and dried. The dot blot was blocked in 5% skim milk in TBST for 1 h. For all blots, portions of the blot were incubated with primary antibodies for 2 h in 1% skim milk in TBST at room temperature (RT). Antibodies used include: anti-candidalysin nanobody (1:5000), 6D11 anti-SLO mAb (1:1000), CPTC-A1–3 anti-annexin A1 mAb (1:250), anti-annexin A2 mAb (1:1000), anti-Alix mAb (1:250), MANLAC-4A7 anti-Lamin A/C mAb (1:250), AC-15 anti-β-actin mAb (1:5000) or with rabbit polyclonal antibodies: anti-aerolysin, anti-annexin A6, anti-MEK, anti-phospho-MEK [Ser217/Ser221], anti-ERK p44/42, anti-phospho-ERK[Thr202/Tyr404], each at 1:1000 dilution. Next, blots were washed 3x for 5 min in TBST and then incubated with HRP-conjugated anti-mouse-IgG, anti-rabbit-IgG, or anti-alpaca V_HH_ antibodies (1:10,000) for 1 h in 1% skim milk in TBST at RT, washed 3x for 5 min in TBST, and developed with enhanced chemiluminescence (ECL): either 0.01% H_2_O_2_ (Walmart, Fayetteville, AR, USA), 0.2 mM p-Coumaric acid (Sigma), 1.25 mM Luminol (Sigma) in 0.1 M Tris-HCl pH 8.4, ECL Plus Western Blotting Substrate (Cat# 32134), ECL Prime Western Blotting Reagent (Cat# RPN2232, Cytiva), or Pierce ECL western blotting substrate (Cat#32106, ThermoFisher Scientific). Full Western blots are included (Supplementary Figs S9-S12).

### Statistics

Prism 10.4.1 (GraphPad, San Diego, CA, USA) or Excel were used for statistical analysis. Sample size was chosen to ensure adequate power to detect the differences in toxin killing. Data are represented as mean ± S.E.M. as indicated. The LC_50_ for toxins was calculated by fitting % specific lysis curve to a logistic model in Excel as previously described^26^. Statistical significance was determined by regular or repeated measures one-way or two-way ANOVA as described. p<0.05 was considered to be statistically significant. Graphs were generated in Excel or Prism.

## Supporting information

Western Blots S9-S12

## Acknowledgments

The authors would like to thank members of the Keyel lab for critical review of the manuscript. We thank colleagues for the generous gifts of reagents. We thank the College of Arts & Sciences Microscopy for use of facilities. This work was supported by American Heart Association grants 16SDG30200001 and 23IPA1053083, Texas Tech University, and NIH grant 1R21AI156225 to PAK. JRN was supported by grants from the Wellcome Trust (214229_Z_18_Z).

## Author Contributions

Conceptualization: RT, VK, CL, BH, JRN, PAK

Methodology: RT, PAK

Investigation: RT, VK, CL, PAK

Data Curation: RT, PAK

Formal Analysis: RT, PAK

Supervision: PAK

Resources: BH, JRN

Project Administration: PAK

Funding Acquisition: PAK, JRN

Writing—original draft: RT, VK, PAK

Writing—review & editing: RT, VK, CL, BH, JRN, PAK

## Conflicts of Interest

PAK is co-founder of Ardiyon Bio. The funders had no role in the design of the study; in the collection, analysis, or interpretation of data; in the writing of the manuscript; nor in the decision to publish the results.

## Data and materials availability

All data are available in the main text or the supplementary materials.

## Supplemental Tables

**Supplementary Table S1.** Lytic parameters of toxins used.

| Toxin | Figure Used | Hemolytic Activity (HU/mL) | Protein Concentration (mg/mL) | Specific Activity (HU/mg) |
| --- | --- | --- | --- | --- |
| Candidalysin WT | 1A, D, E, 2A-C, 3A-E, 4A, B, 5C-I, 6A, S1A, S2A, S3A-D, S4A-E, S5A, S6A, S7A-C | $5 \times 10^4$ | 10 | $5 \times 10^3$ |
| Candidalysin<sup>mutAA</sup> | 1A, 5D, G, 6A, S7A-C | $5 \times 10^3$ | 10 | $5 \times 10^2$ |
| Aerolysin WT | 1B, D, 2A, B, 3A, 4A, C, 5C, D-I, 6A, S3B, C, S4A, S5B, S6B, S7A-C | $4 \times 10^5$ | 2.8 | $1.43 \times 10^5$ |
| | 1E, 2C, 3B-E, 7A, S1B, S2B, S3A, D, S4B-E, | $6.7 \times 10^5$ | 2.4 | $2.8 \times 10^5$ |
| Aerolysin<sup>Y221G</sup> | 1B | $<1 \times 10^3$ | 1.6 | $<6.25 \times 10^2$ |
| SLO WT | 1C-E, 3A, S1C, S2C, S3A, S4A | $6 \times 10^5$ | 1.5 | $4 \times 10^5$ |
| | 2B, 3B, C, 4D, 5C-I, 6A, S3B, S4B, C, S5C, S6C, S7A-C | $1 \times 10^6$ | 2.4 | $4.2 \times 10^5$ |
| | 2A, C, D, 3B-E, 4A, 5C, S3B, D, S4B-E, S6C | $2 \times 10^6$ | 5.3 | $3.8 \times 10^5$ |
| SLO ML | 1C | $<1 \times 10^3$ | 3.2 | $<3.13 \times 10^2$ |

## Supplemental Video Legends

**Video V1. Cells utilize annexin A6 mediated microvesicle shedding to remove candidalysin pores.** HeLa cells transfected with annexin A6-YFP (green) were challenged with sublytic (A) candidalysin (CLY WT), (B) candidalysinAA (CLY mutAA), (C) aerolysin (Aero), or (D) SLO in the presence of 2 μg/mL TO-PRO3 (blue) and imaged at 37°C by live cell confocal imaging at 1-3.5 frame/second. Triton-X-100 was added at the end of 45 min of imaging as a positive control for cell permeabilization. Images were bleach-corrected by histogram matching. Time shows min after toxin addition. Scale bar = 10 μm.

**Video V2. Candidalysin triggers delayed annexin A2 translocation to membrane.** HeLa cells transfected with annexin A2-GFP (green) were challenged with sublytic (A) candidalysin (CLY WT), (B) candidalysinAA (CLY mutAA), (C) aerolysin (Aero), or (D) SLO in the presence of 2 μg/mL TO-PRO3 (blue) and imaged at 37°C by live cell confocal imaging at 1-3.5 frame/second. Triton-X-100 was added at the end of 45 min of imaging as a positive control for cell permeabilization. Images were bleach-corrected by histogram matching. Time shows min after toxin addition. Scale bar = 10 μm.

## Notes

### Summary of Updates

Experiments recapitulating chloride effect in A431 epithelioid cells added. Dot blots to test antibody binding to non-hemolytic Clys added Bilayer experiment added to test direct interaction of ions with Clys Text revisions/updates

