## Supplementary figures and images for "The fungal peptide toxin candidalysin induces distinct membrane repair mechanisms compared to bacterial pore-forming toxins"

### Supplemental Figures S1-S8

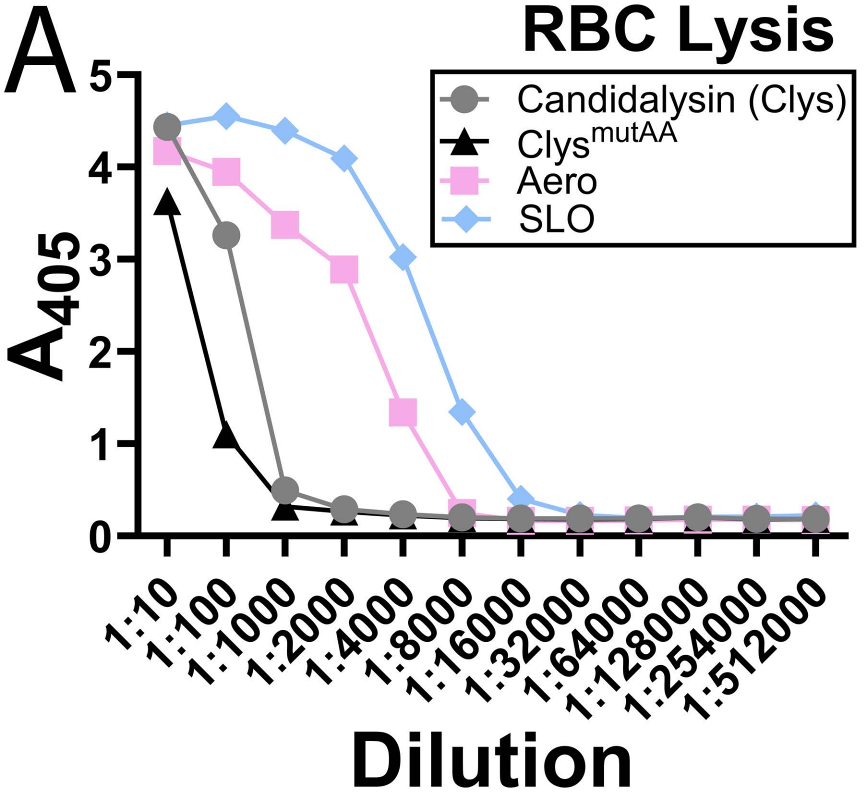

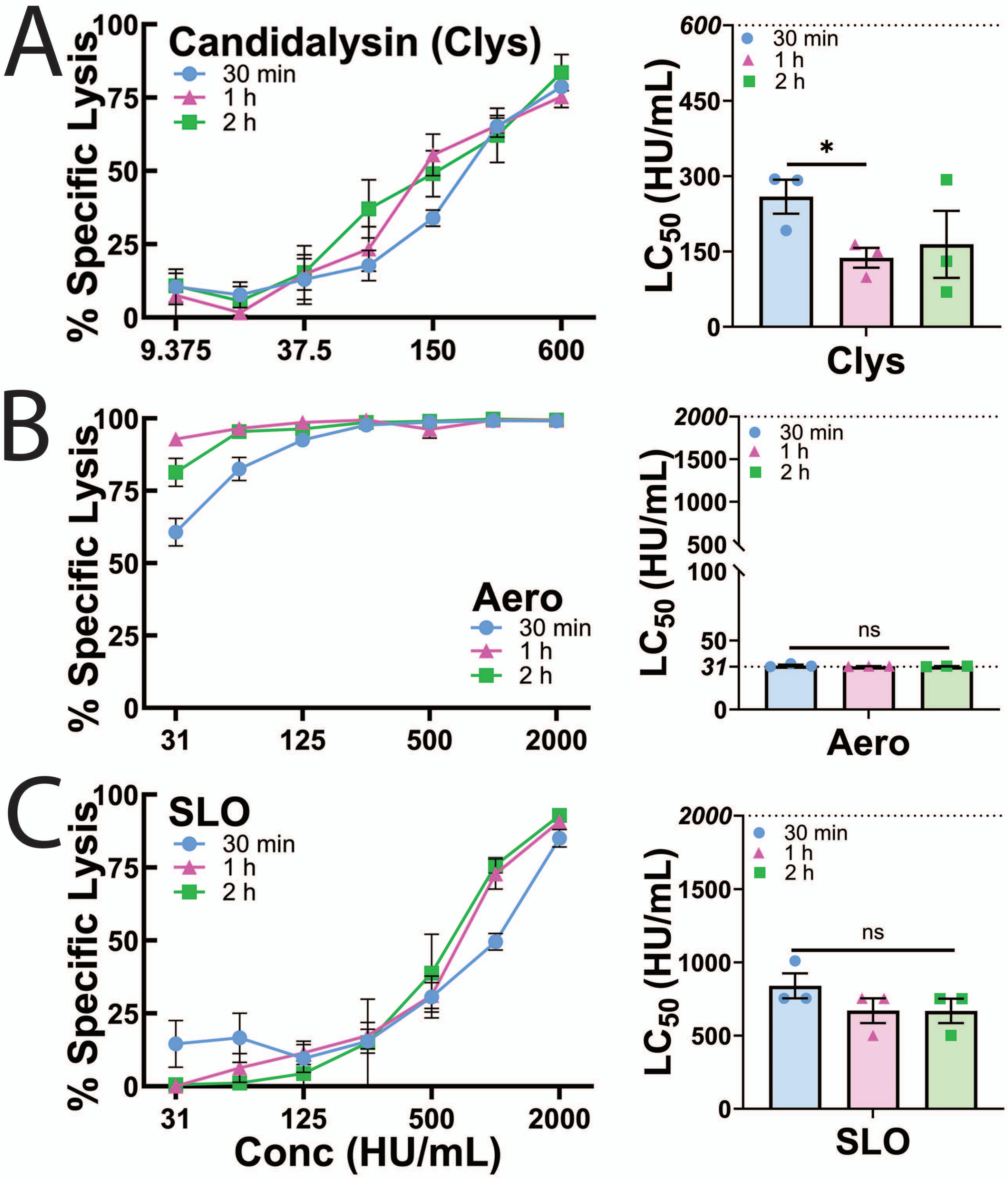

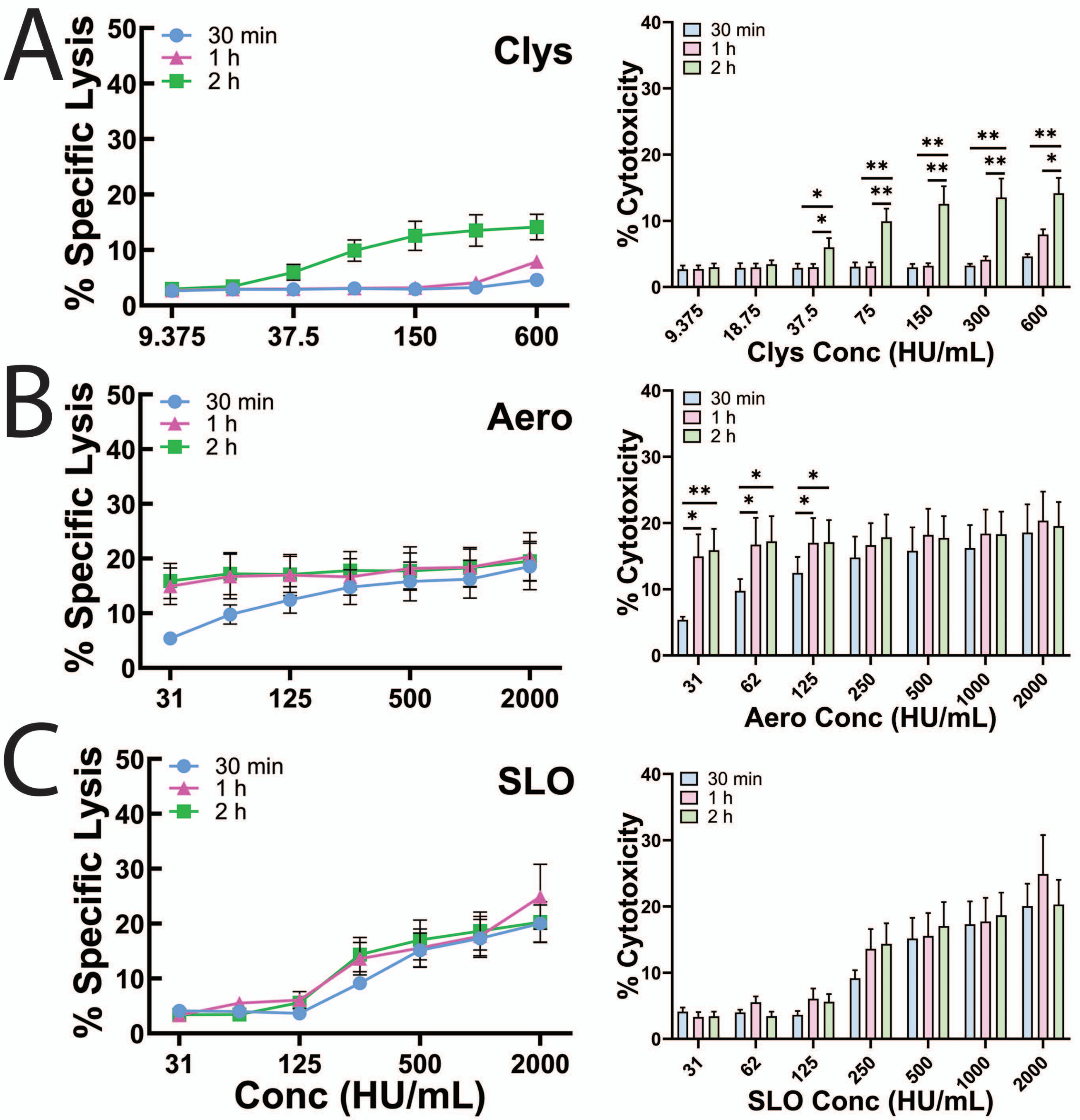

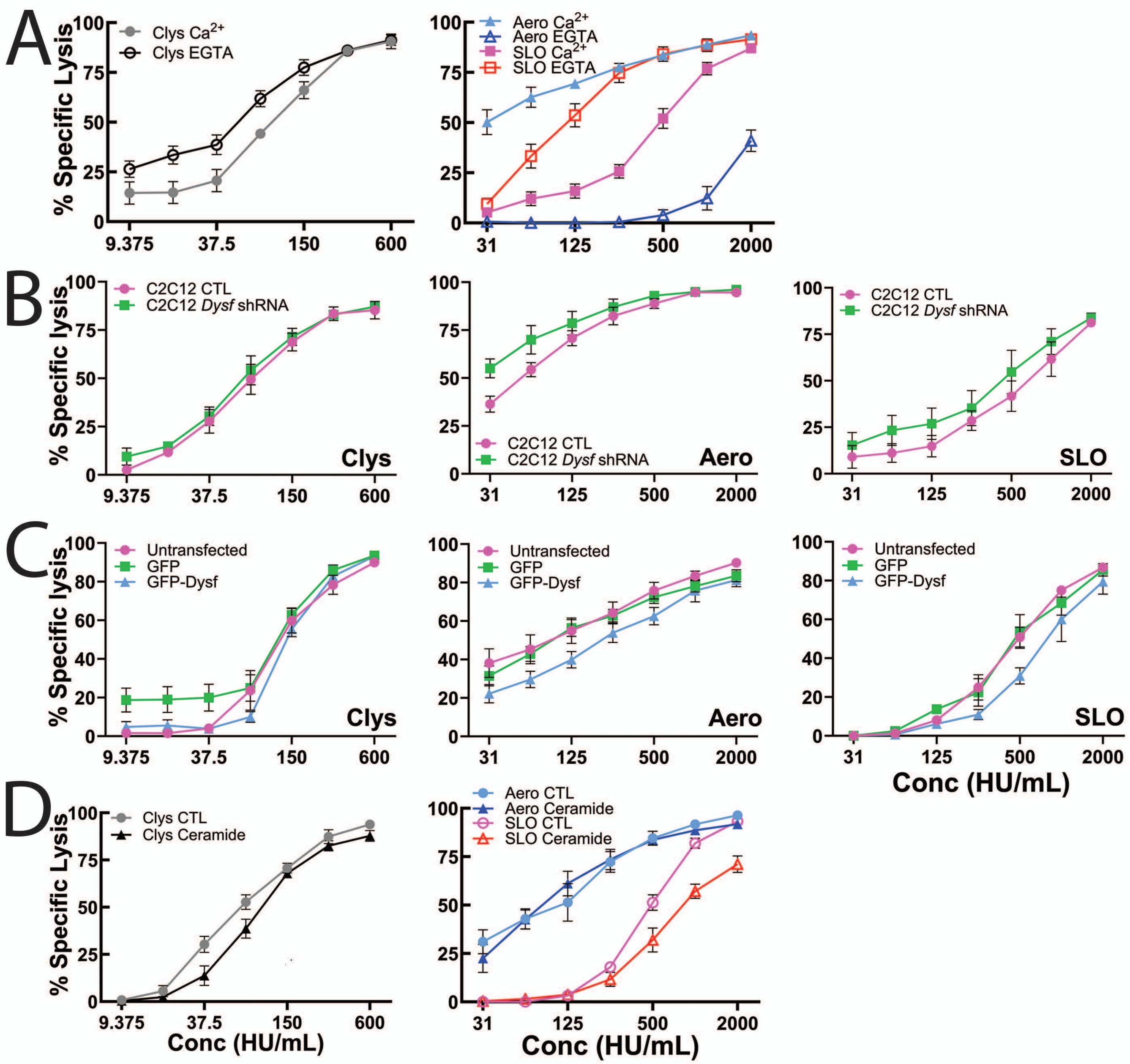

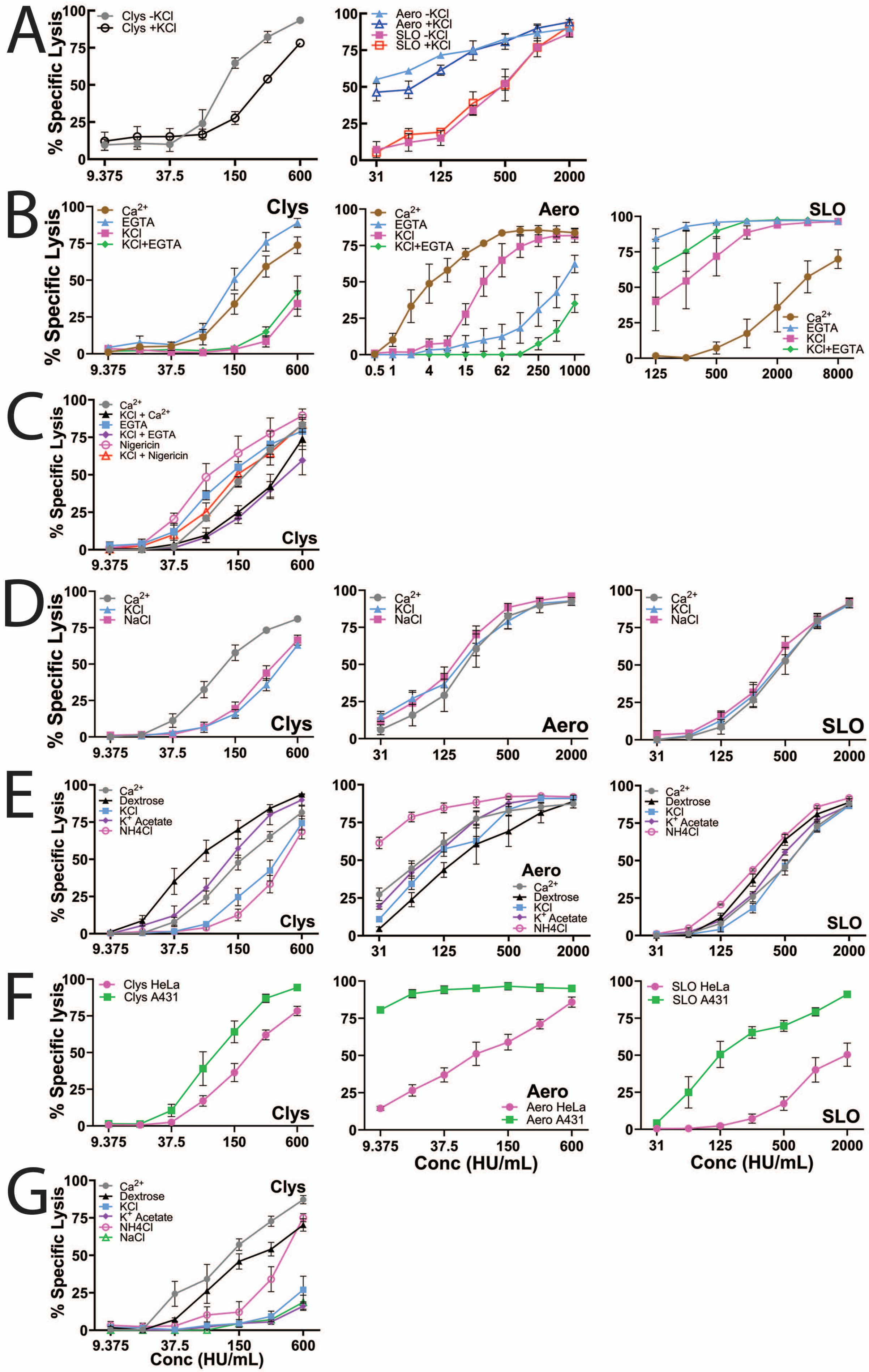

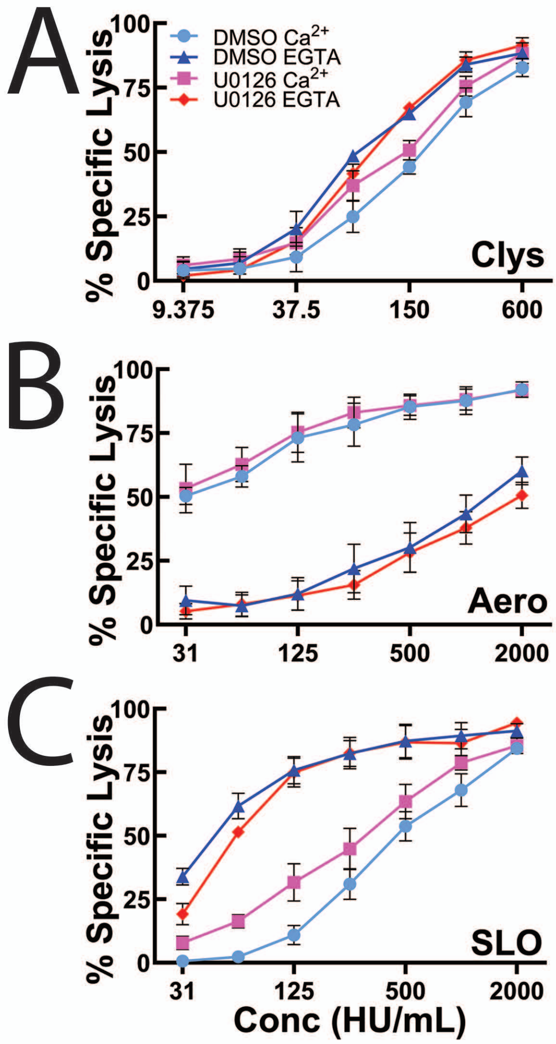

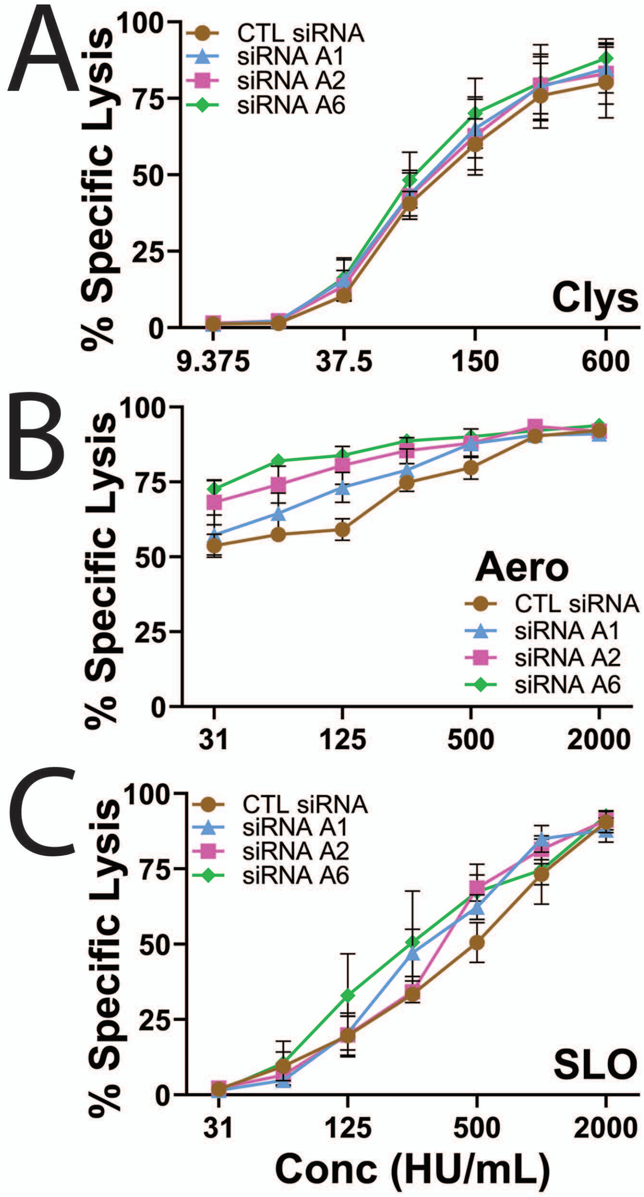

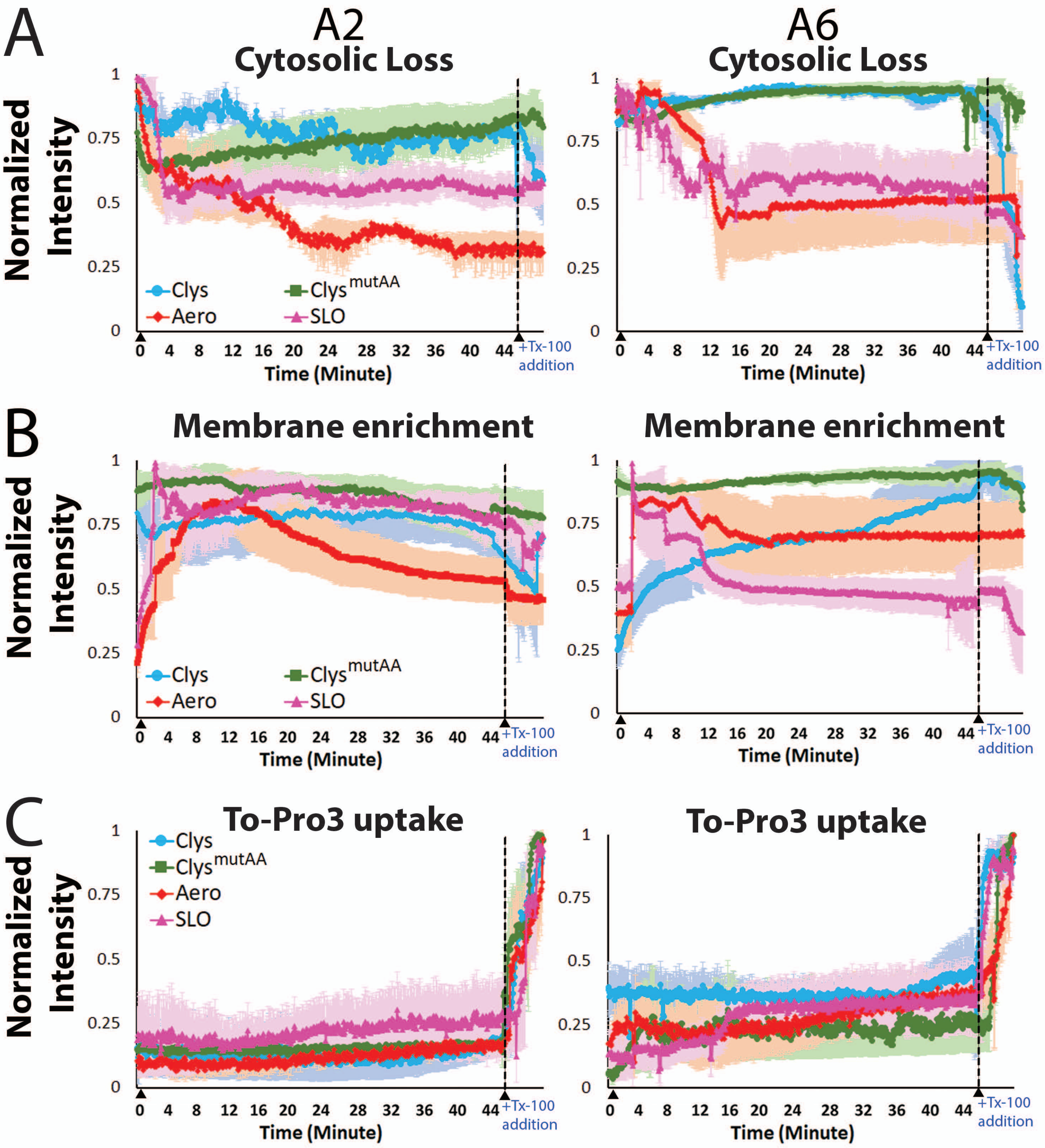

### Western Blots S9-S12

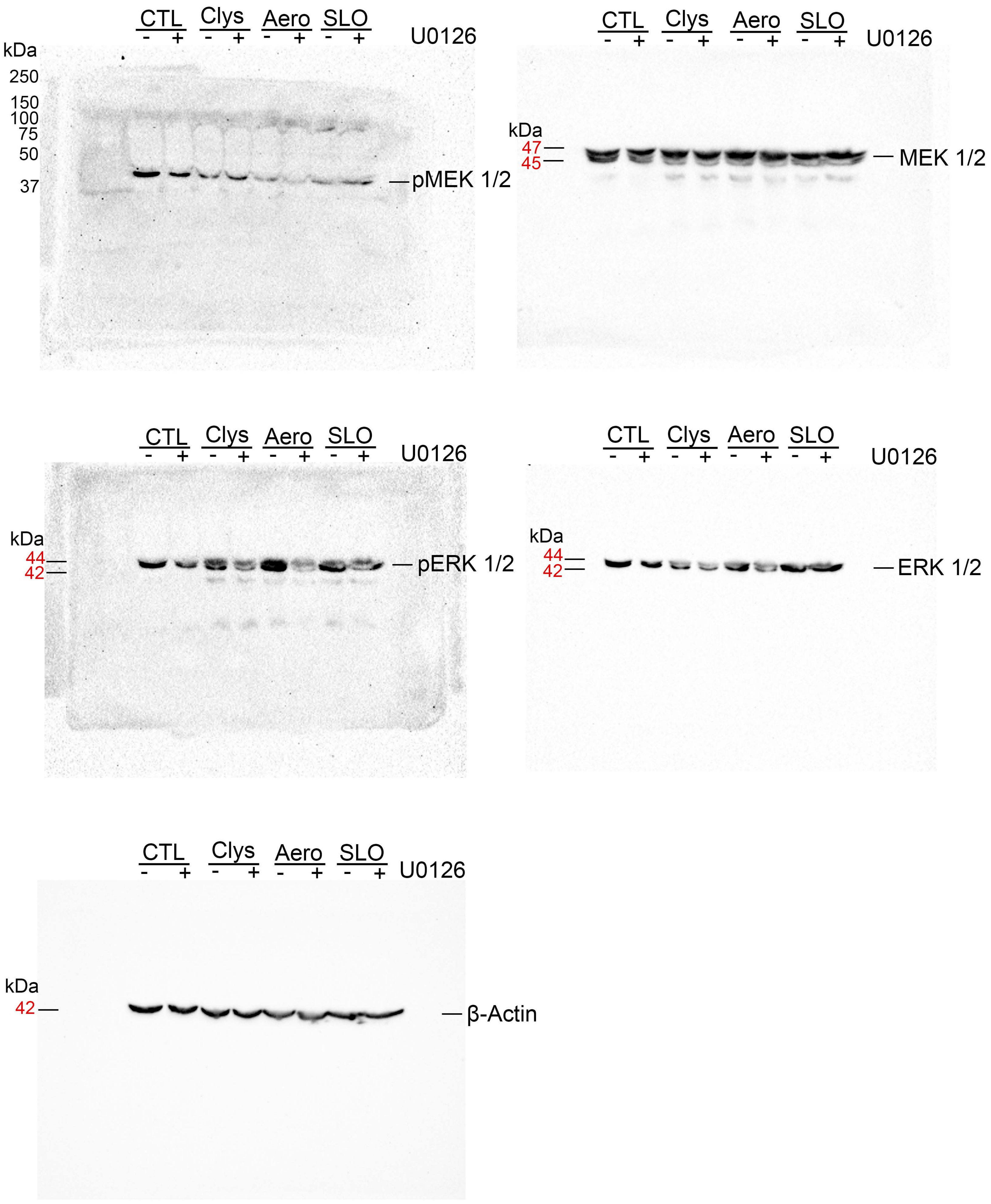

#### Figure S10

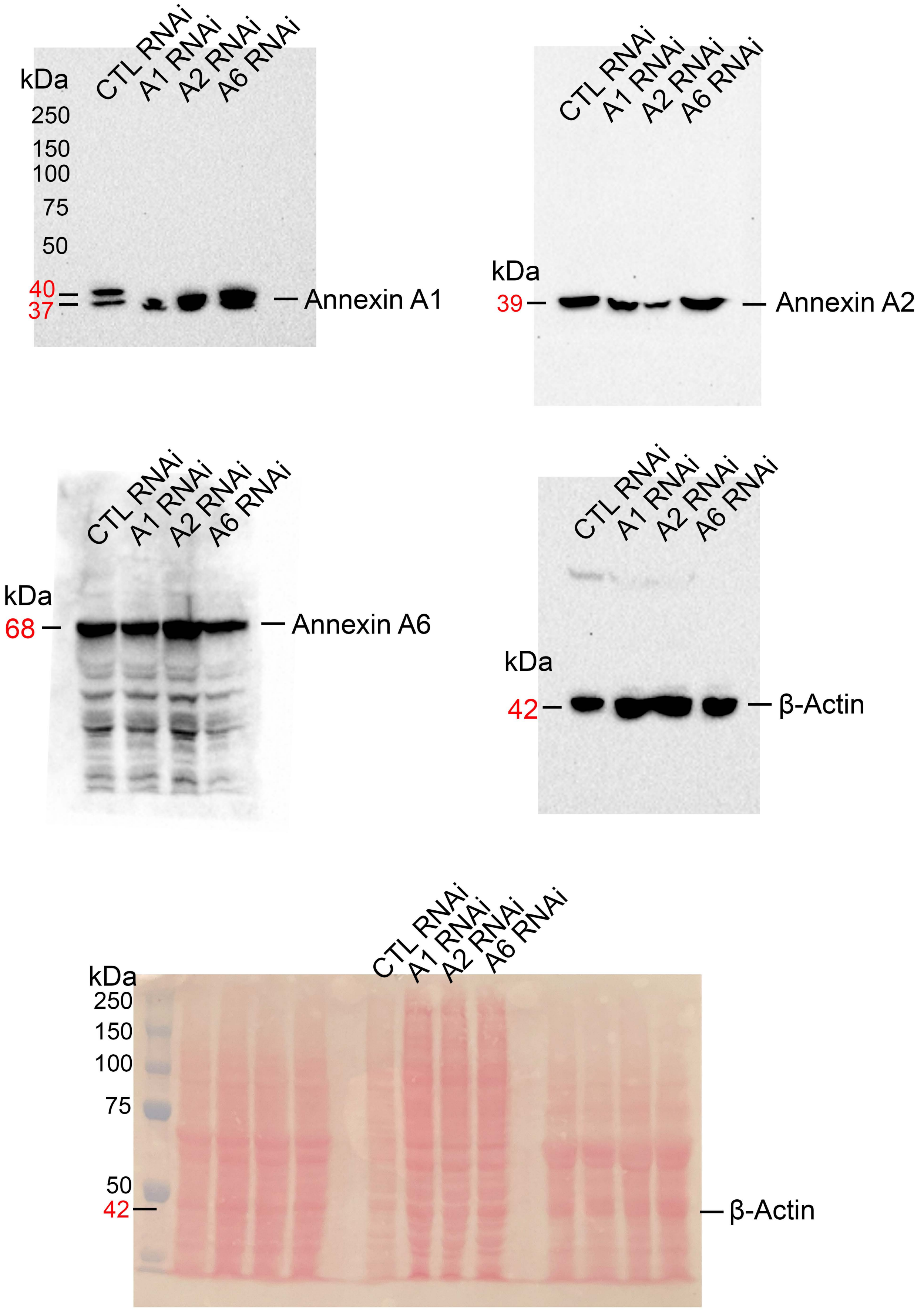


Ponceau stain of the blots

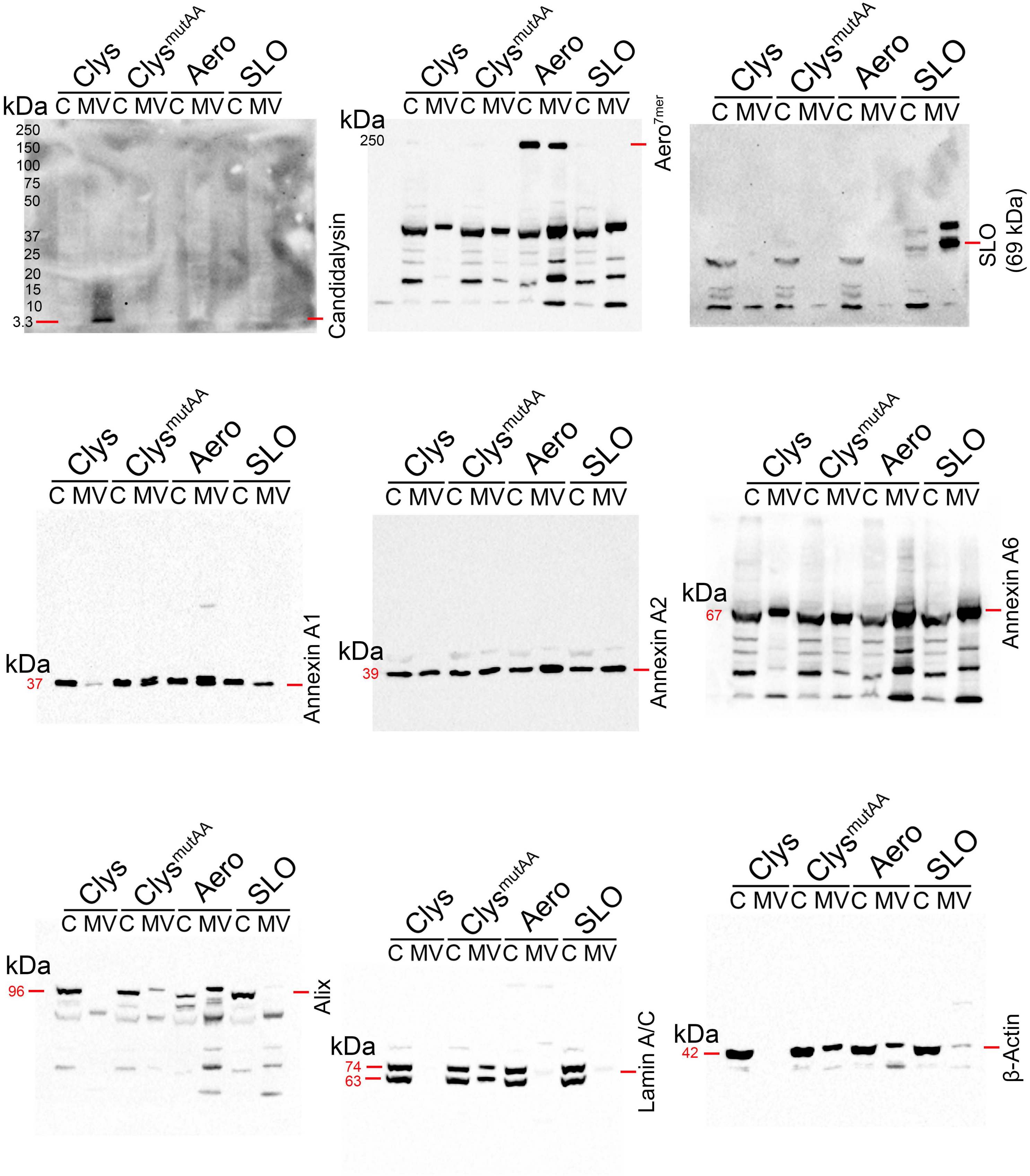

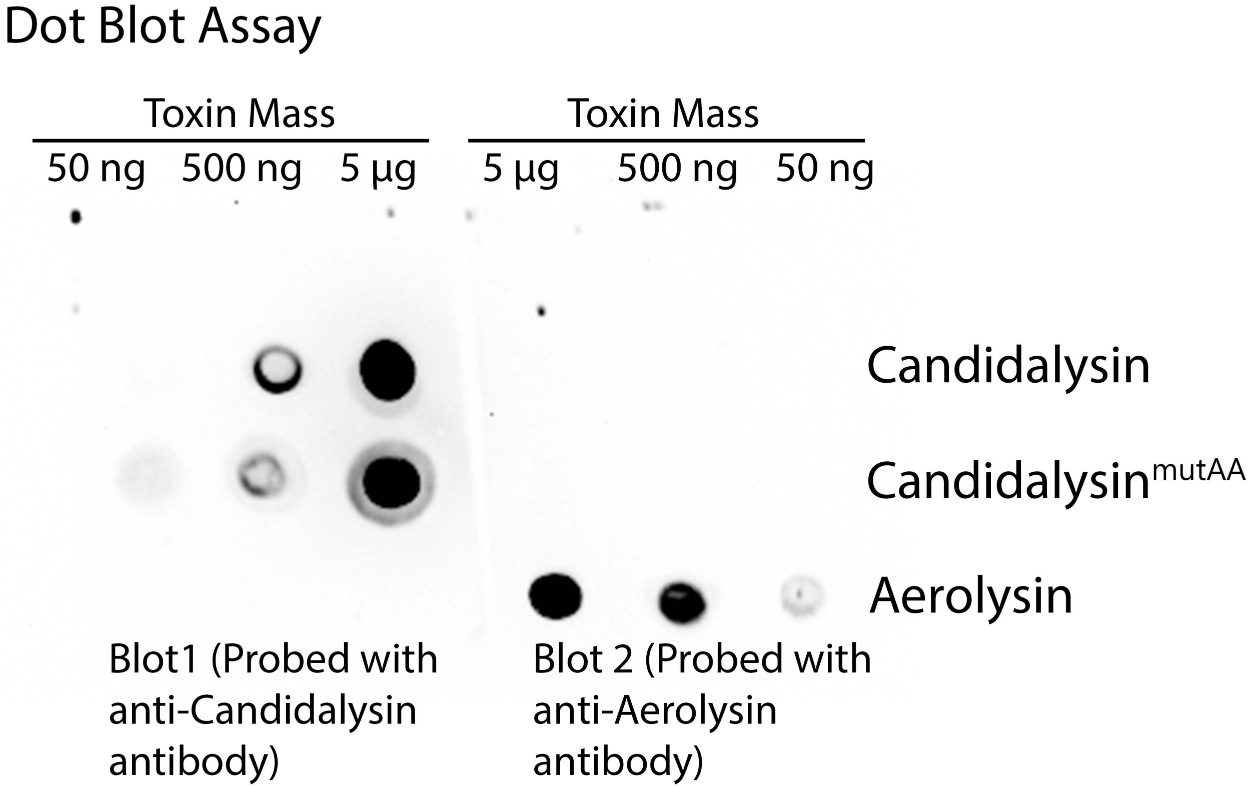
